# Enhanced legumain activity links progranulin deficiency to TDP-43 pathology in frontotemporal lobar degeneration

**DOI:** 10.1101/2024.01.16.575687

**Authors:** Sophie Robinson, Marvin Reich, Maria-Teresa Mühlhofer, Katrin Buschmann, Eline Wauters, Quirin Mühlhofer, Georg Werner, Andrea Ahles, Stefan Engelhardt, Claudia Krenner, Björn Bartels, Simon Gutbier, Anika Reifschneider, Henrick Riemenschneider, Thomas Reinheckel, Dieter Edbauer, Matthew J. Simon, Mihalis S. Kariolis, Todd Logan, Gilbert Di Paolo, Christine Van Broeckhoven, Markus Damme, Dominik Paquet, Christian Haass, Anja Capell

## Abstract

Loss-of-function mutations in *GRN* are a major cause of frontotemporal lobar degeneration (FTLD) with TDP-43-positive inclusions. Progranulin (PGRN) loss leads to lysosomal dysfunction, microglial hyperactivation, and TDP-43 deposition, yet the underlying pathomechanism remains unknown. We demonstrate that PGRN slows the maturation and limits the proteolytic activity of the lysosomal protease legumain (LGMN). Accordingly, LGMN activity is strongly elevated in *Grn* knockout (ko) mice, in human induced pluripotent stem cell-derived *GRN* ko microglia, and in FTLD-*GRN* patients’ brain. Secreted microglial LGMN is internalized by neurons, where it mediates pathological processing of TDP-43, which is prevented by selective LGMN inhibition. In contrast, AAV-mediated overexpression of LGMN in mouse brains promotes TDP-43 processing, the aggregation of phosphorylated TDP-43 and increases plasma neurofilament light chain (NfL), a marker for neuronal loss. Our findings identify LGMN as a link between PGRN haploinsufficiency and TDP-43 pathology in FTLD-*GRN* and suggest LGMN as a therapeutic target.

## Main

Frontotemporal lobar degeneration (FTLD) is a non-curable, rare, early-onset neurodegenerative disorder^1^. Up to 50% of FTLD patients develop TAR DNA binding protein 43 (TDP-43) pathology, which is characterized by aberrant TDP-43 processing, cytosolic deposition, abnormal phosphorylation and nuclear clearance^2,3^. Heterozygous loss-of-function (LOF) mutations in the *GRN* gene are a major cause of FTLD (FTLD*-GRN*) with TDP-43 pathology^4,5^, whereas homozygous LOF mutations result in neuronal ceroid lipofuscinosis (NCL)^6,7^, a lysosomal storage disorder. In the brain, progranulin (PGRN), a secreted growth factor-like protein, is critical for neuronal survival^8^ and diminishes inflammatory processes and hyperactivation of microglia upon CNS damage^9^. As a result, loss of PGRN in mice leads to inflammation and the transition of microglia to a disease associated microglia (DAM) population^10-12^. Microglial dysregulation upon PGRN loss may be associated with its function within lysosomes^13^. NCL cases caused by loss of PGRN and PGRN-deficient mouse models exhibit multiple endo-lysosomal and autophagic phenotypes, such as dysregulation of lysosomal proteases and pH^14-20^, lipid dyshomeostasis, and lipofuscin accumulation^13,16,21-25^. PGRN is either secreted or transported to lysosomes *via* multiple pathways, including sortilin and prosaposin/cation-independent mannose-6- phosphate receptor^26^. Upon increased acidification of the endo-lysosomal pathway, PGRN is proteolytically processed into granulin peptides by the lysosomal protease legumain (LGMN), also known as asparagine endopeptidase (AEP), and probably by various cathepsins, including cathepsin B and L^27-29^. An essential role of granulins in lysosomes is suggested, since lysosomal dysfunction caused by PGRN deficiency is ameliorated by granulin peptides^30^. Despite intense research, both the lysosomal function of PGRN / granulins and the mechanistic link between PGRN deficiency and TDP-43 aggregation remain unclear. A spatial paradox, namely PGRN expression primarily in microglia and TDP-43 pathology in neurons, makes the understanding of the link between PGRN loss and TDP-43 pathology even more challenging.

We have previously shown that loss of PGRN causes lysosomal dysfunction with altered cathepsin maturation^18^. To establish a functional link between PGRN deficiency and TDP-43 pathology, we hypothesize that lysosomal dysfunction contributes to disease-associated proteolytic TDP-43 processing. We therefore searched for a protease which plays a key role in processing of TDP-43 and lysosomal cathepsins. LGMN, a caspase-like cysteine protease of the C13 peptidase family emerged as a prime candidate^31^, since it is involved in the proteolytic turnover of various cathepsins^32,33^ and capable of cleaving TDP-43 *in vitro*^34^. Here, we provide a mechanistic link between PGRN deficiency and TDP-43 pathology *via* dysregulation of LGMN.

## Results

### PGRN deficiency results in elevated LGMN maturation and activity

To test our hypothesis, we investigated LGMN expression and activity in various model systems and in brain material from FTLD-*GRN* patients. Mature LGMN and its proteolytic activity were strongly elevated in *Grn* ko MEF compared to wt MEF (Fig. 1a,b). In line with that, knockdown of LGMN blunts the previously described increased maturation and accumulation of the heavy chain of cathepsins D and L in the *Grn* ko MEF (Extended Data Fig. 1a–c)^18^. *LGMN* ko mice further proved that LGMN is required for the generation of the heavy chain form of cathepsin L^35^ and cathepsin D in mice (Extended Data Fig. 1d). Moreover, elevated levels of mature LGMN were detected in the brain tissue of 6-month-old *Grn* ko mice (Fig. 1c) and increased proteolytic activity of LGMN was observed in all ages investigated between 3 and 24 months (Fig. 1d), indicating that LGMN dysfunction might be a driver of the disease and not a response. Increased LGMN expression and activity were not a result of transcriptional changes (Fig. 1e), suggesting posttranscriptional regulatory mechanisms. To test whether the increase in LGMN maturation and activity is directly caused by the loss of PGRN, we reintroduced PGRN in *Grn* ko MEF and *Grn* ko mice (Extended Data Fig. 2a-e). Restoration of PGRN in *Grn* ko MEF by stable transfection reduced mature LGMN and completely normalized LGMN hyperactivity (Extended Data Fig. 2a-b). To reintroduce PGRN *in vivo*, we used the previously described protein transport vehicle (PTV):PGRN, which shows enhanced brain penetrance due to its ability to bind to the transferrin receptor (TfR), to restore brain PGRN levels in *Grn* ko mice expressing chimeric humanized TfR (*Grn* ko; *TfR* ki) (Extended Data Fig. 2c)^16^. PTV:PGRN reduced maturation and activity of LGMN over time (Extended Data Fig. 2d-e). Thus, PGRN appears to directly affect LGMN maturation and consequently its proteolytic activity.

**Fig. 1:**
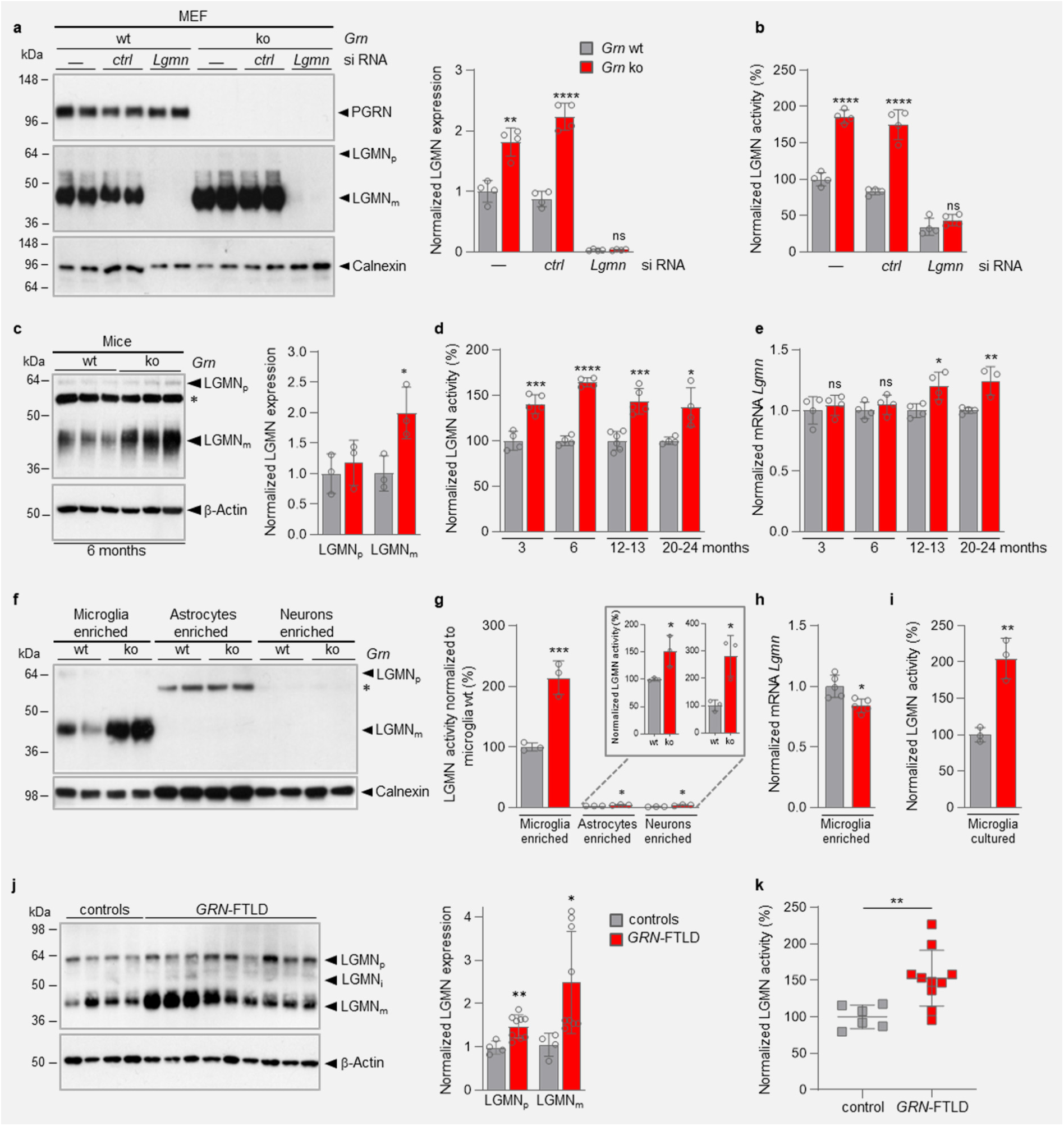
PGRN deficiency leads to enhanced LGMN maturation and elevated activity. **a**, Representative immunoblots of mouse embryonic fibroblasts (MEF), isolated from wild-type (wt) and granulin knockout (*Grn* ko) mice, probed for progranulin (PGRN), legumain (LGMN), and calnexin as loading control. MEF are either non-(-), control-(*ctrl*) or *Lgmn* siRNA-transfected. Quantification of LGMN expression normalized to non-transfected wt levels (n=4). **b**, *In vitro* LGMN activity of non-(-), mock, *ctrl* and *Lgmn* siRNA-transfected *Grn* ko MEF normalized to non-transfected wt MEF (n=4). **c**, Representative immunoblot for LGMN in total brain homogenates of 6-month-old wt and *Grn* ko mice. Proform (LGMN_p_), mature form (LGMN_m_), and non-specific band (*) are indicated, β-actin verified equal loading. Quantification of the immunoblot signals normalized to wt (n=3 mice per genotype). **d**, *In vitro* LGMN activity of brain homogenates from 3-, 6-, 12-13-, and 20-24-month-old wt and *Grn* ko mice (n=4-6 per genotype and age group). **e**, Total brain mRNA of 3-, 6-, 12-13-, and 20-24-month-old mice normalized to wt (n=4-5). **f**, Representative immunoblot for LGMN in MACS-sorted microglia-, astrocyte- and neuron-enriched brain cell fractions of 4-5-month-old wt and *Grn* ko mice. LGMN_p_ and LGMN_m_ are indicated, calnexin verified equal loading for each fraction. **g**, Microglia-, astrocyte- and neuron-enriched fractions from 4-5-month-old *Grn* ko and wt mice analyzed for LGMN activity either normalized to wt microglia or wt of the respective cell type (insert) (n=3 mice per cell type and genotype). **h**, *Lgmn* mRNA levels of microglia isolated from 6-month-old wt and *Grn* ko mice normalized to wt (n=5). **i,** LGMN *in vitro* activity in lysates of cultured primary mouse microglia (n=3 mice per genotype). **j**, **k**, LGMN expression (**j**) and activity (**k**) analyzed in brain lysates of frontal cortex from FTLD/*GRN* patients and pathology-negative control cases (see Extended Data Table 1). LGMN_p_ and LGMN_m_ are quantified in the immunoblot and normalized to the mean signal of control cases (**j**). Data are mean ± s.d. of biologically independent experiments; unpaired two-tailed t-test: ns not significant; \**P*<0.05, \*\**P*<0.01, \*\*\**P*<0.001, \*\*\*\**P*<0.0001. *P* values and statistical source data are provided.

In the brain, PGRN is predominantly expressed by microglia^11,18^. In contrast, pathological TDP-43 inclusions observed in FTLD or amyotrophic lateral sclerosis (ALS) are mainly localized in neurons and are less abundant in glial cells^2,3^. Thus, we next investigated cell type-specific expression of LGMN. LGMN and PGRN were both strongly enriched in acutely isolated microglia (Fig. 1f; Extended Data Fig. 3). Moreover, microglia showed efficient LGMN maturation as well as the highest LGMN activity of all cell types investigated (Fig. 1f-g; Extended Data Fig. 3). Even though LGMN expression in neurons and astrocytes was much lower than in microglia, the relative change in LGMN activity between PGRN deficient and wt cells was highest in neurons with a 3-fold increase in *Grn* ko (Fig. 1g). In contrast to the LGMN activity, *Lgmn* mRNA levels were not increased in acutely isolated microglia (Fig. 1h), supporting the conclusion that LGMN is regulated at a posttranscriptional level. In addition, in primary cultures of *Grn* ko microglia LGMN activity was twice as high as in wt controls, mirroring the results obtained from the acutely isolated microglia (Fig. 1i). To investigate whether elevated LGMN levels are disease relevant, we assessed LGMN maturation and activity in the frontal cortex (Brodmann area 9 and 10) of patients suffering from FTLD-*GRN* (Extended Data Table 1). In line with our findings in the model systems described above, we observed increased levels, maturation and proteolytic activity of LGMN in patient brains relative to age matched non-diseased controls (Fig. 1j,k).

### PGRN slows LGMN auto-activation

Next, we investigated whether PGRN directly affects LGMN catalytic activity or interferes with its proteolytic activation. With decreasing pH during lysosome maturation, LGMN is autoproteolytically activated in multiple steps and processed to its final mature form of approximately 43 kDa by unknown proteases (Fig. 2a)^36^. Autocatalytic *in vitro* maturation of the pro-form of LGMN (pro-LGMN) to an intermediate active form has been previously shown to be facilitated at an acidic pH^31^. To investigate whether PGRN directly affects LGMN activity, we measured the *in vitro* activity of autocatalytically pre-activated recombinant (r) LGMN in the presence and absence of rPGRN. PGRN did not affect the catalytic activity of rLGMN (Fig. 2b). Based on the increased LGMN maturation in various PGRN deficient model systems (Fig. 1), we next investigated whether the presence of rPGRN affects maturation of rLGMN.

**Fig. 2:**
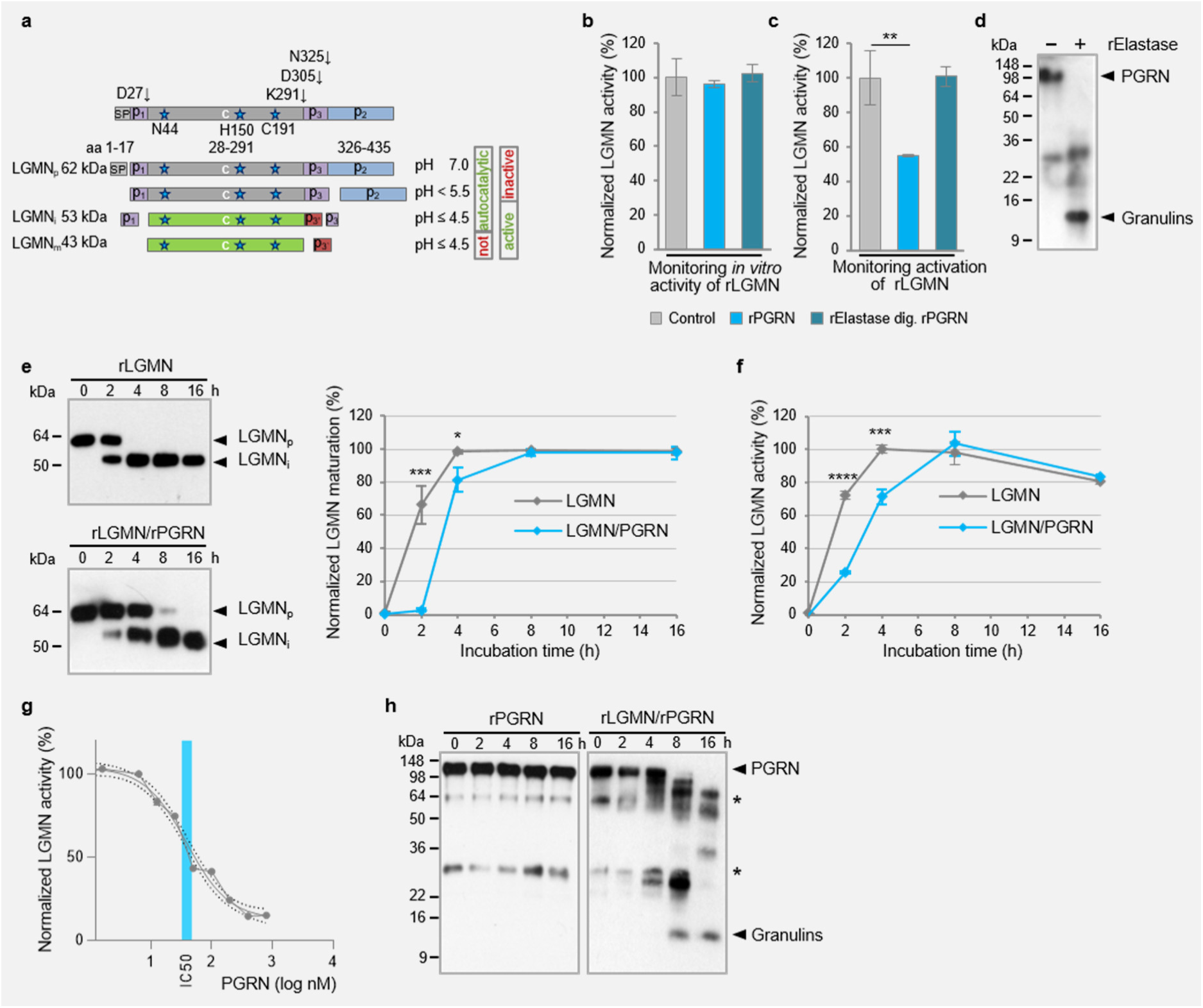
PGRN modulates LGMN activation and is proteolytically processed by LGMN. **a**, Schematic representation of pH-dependent auto- and not auto-catalytic LGMN maturation. N- and C-terminal pro-peptides (p1-p3) and the mature chain (c) with the active site residues (blue stars) are specified. Proform (LGMN_p_), intermediate-form (LGMN_i_) and mature-form (LGMN_m_) are indicated. **b**, **c**, Activity analysis of *in vitro* auto-catalytically matured recombinant (r) LGMN incubated for 4 h at acidic pH. Either rPGRN or elastase-digested PGRN were added after (**b**) or before the 4 h activation step (**c**) (n=3). **d**, PGRN digest with elastase controlled by immunoblotting. **e**, **f,** Activation of rLGMN with/without rPGRN at acidic pH for indicated time points, monitored by immunoblotting. Quantification of the relative LGMN_i_ level (% of total LGMN) (**e**) or activity (**f**) for each timepoint (n=3). **g**, LGMN activity after 2 h incubation with different concentrations of rPGRN at pH 4. The IC_50_ was calculated using a non-linear curve fit, 95% confidence intervals (CI) are indicated (n=3). **h**, Immunoblot analysis of rPGRN turnover and granulin peptide generation by rLGMN. Stars indicate rLGMN independent bands. Data are mean ± s.d. of independent technical replicates, unpaired two-tailed t-test: ns, Strikingly, rLGMN activity was reduced by more than 50 % when rPGRN was added to pro-rLGMN during the activation step at acidic pH 4 (Fig. 2c), while elastase-digested rPGRN did not reduce rLGMN activation (Fig. 2c,d), suggesting that rPGRN, but not granulins, regulate the maturation of rLGMN. Autocatalytic maturation of pro-rLGMN was then monitored longitudinally at pH 4.0 with and without rPGRN. Proteolytic *in vitro* maturation and enzymatic activation of rLGMN under acidic conditions was significantly slowed in the presence of rPGRN (Fig. 2e,f). Moreover, rPGRN reduced autocatalytic activation of rLGMN in a dose-dependent manner with an IC50 of 39.28 nM (Fig. 2g). However, after full autocatalytic maturation (Fig. 2e), the catalytic activity of rLGMN was identical in the presence and absence of rPGRN (Fig. 2f). Thus, PGRN does not affect the catalytic activity of LGMN *per se*, but rather slows its autocatalytic maturation. Intriguingly, as suggested before^29^, rLGMN efficiently generates granulin peptides *in vitro* (Fig. 2h). After 8 h of co-incubation of rPGRN with rLGMN, full length rPGRN was completely converted into single or multiple granulin peptides. At this time point also full conversion of rLGMN to the intermediate form was observed (Fig. 2e), further indicating that full length PGRN is needed to slow LGMN activation and that LGMN regulates its own maturation by processing of PGRN in a negative feedback loop. *In vivo*, the interaction of LGMN with PGRN is supported by reduced generation of granulins in *Lgmn* ko mice. Granulin peptides are readily detected within purified liver lysosomes of wt mice, but are substantially reduced in *Lgmn* ko mice (Extended Data Fig. 4a,b). Thus, LGMN regulates its own maturation by processing of PGRN in a negative feedback loop.

### Increased LGMN activity promotes pathological processing of TDP-43

Since TDP-43 is known to undergo pathological proteolytic processing^2,37,38^ and contains several potential cleavage sites for LGMN^34^ (Fig. 3a), we investigated if the lack of PGRN affects processing of TDP-43 *via* enhanced LGMN activity. In *Grn* ko MEF we observed a reduction of full-length TDP-43 accompanied by the enhanced formation of TDP-43 fragments (TDP-43F) which were enriched in the urea solubilized fraction (Fig. 3b). Furthermore, C-terminal fragments of TDP-43 (TDP-43CTF) with an approximate molecular weight of 25 kDa, reminiscent of those observed in FTLD cases^34^, were observed in lysates of *Grn* ko MEF (Fig. 3b,c). Abnormal proteolytic processing of TDP-43 was completely blocked upon knockdown of LGMN in *Grn* ko MEF (Fig. 3b,c), proving that disease-defining TDP-43 fragments are generated in a LGMN-dependent manner. Moreover, even in the presence of PGRN, reduction of LGMN was sufficient to reduce TDP-43 processing (Fig. 3b). Immunofluorescence staining of TDP-43 revealed significant cytoplasmic mis-localization of TDP-43 in *Grn* ko MEF (Fig. 3d, upper panels). MEF *Grn* ko show enhanced co-localization of TDP- 43 with LGMN- and LAMP1-positive compartments, indicating that TDP-43 processing by LGMN occurs in the lysosome (Fig. 3d, lower panels).

**Fig. 3:**
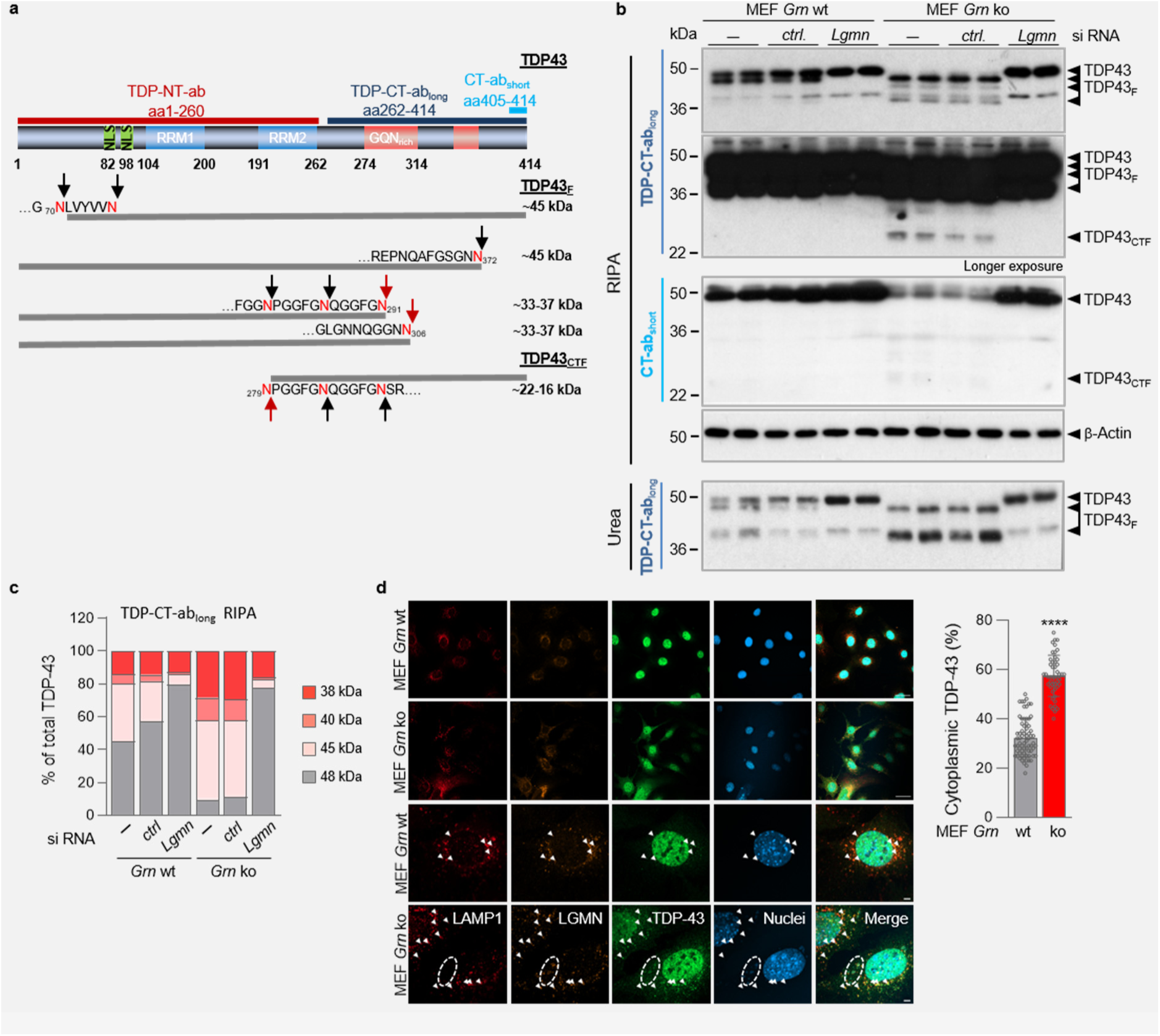
Enhanced LGMN activity results in accelerated TDP-43 processing. **a**, Schematic representation of the TDP-43 protein indicates the nuclear localization sequence (NLS, green), the RNA-recognition motifs (RRM1 / 2, blue), and the G-, Q-, N-rich hnRNP-interacting domains (red). Epitopes of different N-terminal (NT) and C-terminal (CT) antibodies are indicated. Confirmed LGMN cleavage sites (black arrows) and putative fragments are indicated below. Red arrows indicate LGMN cleaved fragments identified in patient material (Herskowitz et al, 2012; Kametani et al, 2016). **b**, Immunoblot analysis of TDP-43, TDP-43 fragments (TDP43F) and C-terminal fragments (TDP43CTF) in MEF *Grn* wt and ko with and without *Lgmn* siRNA-mediated knockdown. **c**, Quantification of TDP-43 processing; holoprotein (48 kDa) and fragments (45 kDa, 40 kDa and 38 kDa) were normalized to total TDP-43 (n=4). **d**, Immunofluorescence staining of the lysosomal protein LAMP1 (deep red), LGMN (orange), TDP-43 (green) and nuclear DAPI staining (blue) of *Grn* wt and ko MEF (top panels scale-bar 10 μm, lower panels 20 μm). Arrowheads indicate co-localization of LAMP1, LGMN and TDP-43, encircled is a cytoplasmic DAPI, LGMN and TDP-43 positive area. Quantification of % cytoplasmic vs. total TDP-43. Data are mean ± s.d. (n=48-69 cells / genotype from 6-8 images), unpaired two-tailed t-test, \*\*\*\**P*<0.0001. *P* values and statistical source data are provided

Taken together, our findings suggest that reduced PGRN results in enhanced LGMN activation, which in turn mediates pathological processing and mis-localization of TDP- 43.

### Shuttling of LGMN from microglia to neurons

TDP-43 deposits are predominantly found in neurons, however compared to microglia, neurons express substantially less PGRN and LGMN and therefore show much less LGMN activity (Fig. 1f,g, Extended Data Fig. 3). Since pro-LGMN was found to be secreted in peripheral immune cells^36^, we hypothesized that PGRN-deficient hyperactivated microglia also release pro-LGMN, which could be internalized by neurons, where it then mediates pathological TDP-43 processing. To provide evidence that catalytically inactive pro-LGMN can be taken up and activated by neurons, we incubated primary mouse hippocampal neurons with conditioned media from wt, inactive mutant (mt) LGMN-, or mock-transfected HeLa cells (Fig. 4a). Robust levels of overexpressed wt and mt LGMN were detected in HeLa cells. Autocatalytic processing occurred in wt, but was altered in mt LGMN expressing cells, suggesting bimolecular processing by endogenous LGMN or alternative processing (Fig. 4b)^39,40^. Upon transfection, conditioned media contained significant amounts of secreted wt or mt pro-LGMN (Fig. 4c). Only secreted wt LGMN was autocatalytically activated upon incubation under acidic conditions while no activity was observed in conditioned media of mt LGMN or non-transfected HeLa cells (Fig. 4d). Both, wt and mt LGMN were internalized and either autocatalytically or alternatively processed in primary neurons (Fig. 4e). Noticeably, processed mt LGMN accumulated in lysates (Fig. 4e). Only neurons incubated with wt LGMN-containing conditioned media show increased LGMN activity (Fig. 4f). Strikingly, we observed enhanced TDP-43 processing relative to total TDP-43 levels and even the formation of pathological TDP-43CTF upon uptake of wt LGMN, but not of the catalytically inactive variant (Fig. 4g). This demonstrates that proteolytically active LGMN is sufficient to generate disease-relevant TDP-43 fragments and that neurons can internalize and activate LGMN.

**Fig. 4:**
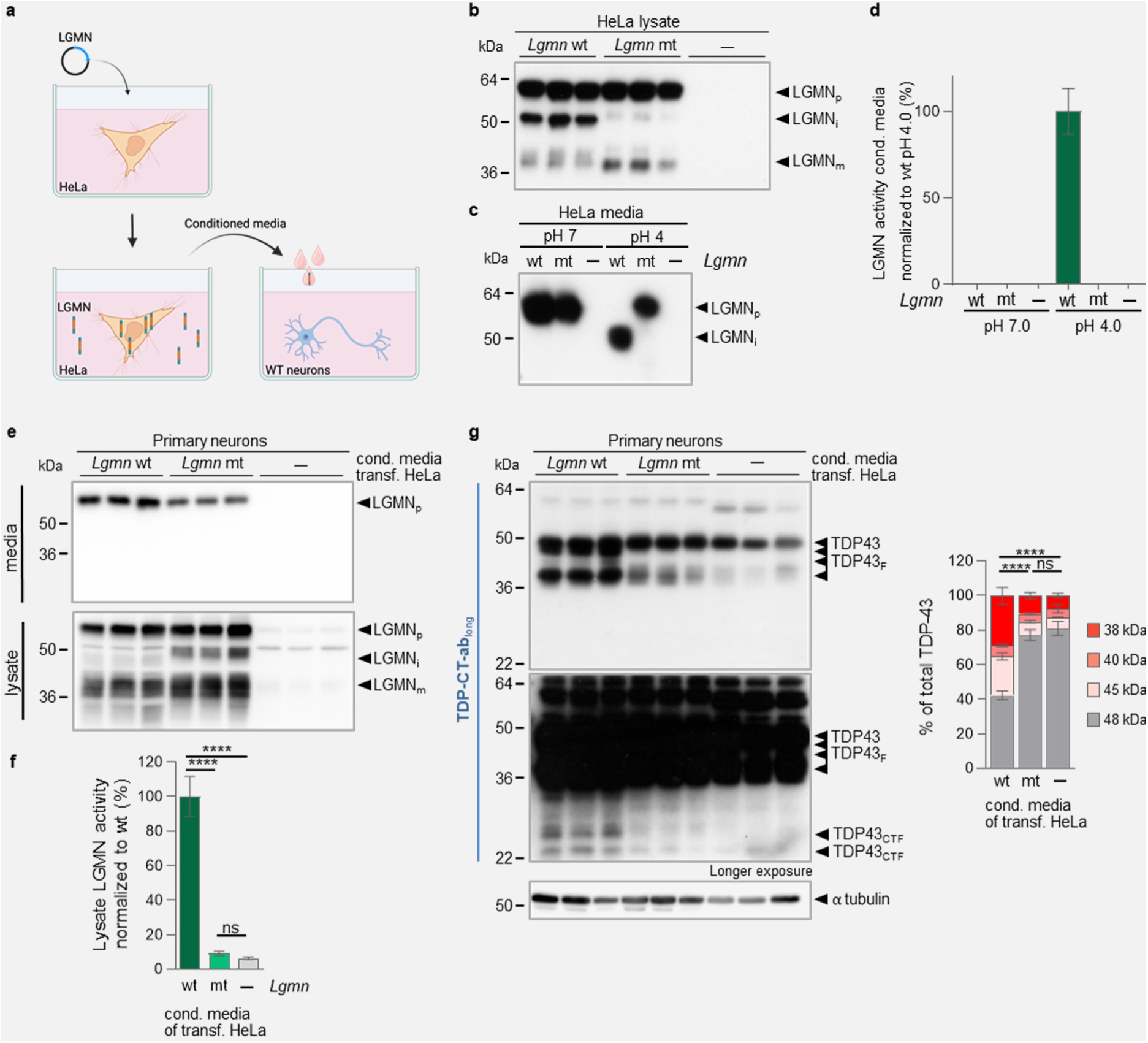
LGMN internalized by neurons results in accelerated TDP-43 processing. **a**, Schematic representation of the media transfer experiment. HeLa cells were transfected with wt mouse LGMN, the proteolytically inactive LGMN C191A variant (mt) or mock (-) and 25% conditioned media supplemented to primary hippocampal mouse neurons for 24 h. (**b,c)**, LGMN expression (LGMN_p_, LGMN_i_, and LGMN_m_) (**b**) and secretion of LGMN_p_ (**c**) by HeLa cells was verified by immunoblotting (n=3 independent experiments). (**c,d**) To monitor autocatalytic processing of LGMN, conditioned media was incubated at pH 7 or pH 4 for 4 h. **d**, LGMN activity of conditioned media incubated at pH 7 and pH 4 (n=3 independent experiments). (**e-g)**, Primary neurons were analyzed after 24 h incubation with conditioned media. Presence of the LGMN proform was confirmed in culture media (**e**). LGMN uptake by neurons (**e**) and TDP-To provide evidence for a pathological link between PGRN reduction, LGMN activation, and enhanced TDP-43 processing in a human disease-relevant system, we generated human iPSC lacking PGRN (*GRN* ko iPSC)^22^ and differentiated them into human induced pluripotent stem cell-derived microglia (hiMGL) and neurons (hiNE). *GRN* ko hiMGL showed a 3-4-fold increased expression and activity of LGMN compared to wt hiMGL (Fig. 5a,b) but only a 1.5-fold increase of *LGMN* mRNA levels (Fig. 5c). Thus, the difference in LGMN mRNA levels between wt and *GRN* ko hiMGL was less pronounced than on LGMN protein or activity levels, further supporting a post-transcriptional regulation of LGMN activity upon PGRN deficiency. In line with our findings in mice (Fig. 1f,g), hiNE had almost no LGMN protein expression or activity (Fig. 5a,b) and showed lower transcript levels than hiMGL (Fig. 5c). Furthermore, *GRN* ko hiNE did not show an increase of LGMN activity in contrast to *Grn* ko mouse neurons (Fig. 1g). To investigate a potential cross-talk between microglia and neurons upon PGRN deficiency, we co-cultured *GRN* wt hiNE with either *GRN* wt or ko hiMGL (Fig. 5d). In co-cultures of wt hiNE with GRN ko hiMGL, we observed increased mature and active LGMN in lysates (Fig. 5f,g). As a consequence, co-culture of *GRN* ko hiMGL with *GRN* wt hiNE showed enhanced TDP-43 processing, indicated by reduced TDP- 43 full length and enhanced TDP-43 fragments (Fig. 5h). To investigate if TDP-43 processing in the co-culture occurs predominantly within neurons, we selectively blocked neuronal LGMN activity by AAV9 (AAV-hSyn-Cst7) mediated neuronal expression of cystatin F (CSTF) prior to co-culturing with hiMGL (Fig. 5d). CSTF and CSTC are brain expressed^41,42^ effective inhibitors of LGMN (Extended Data Fig. 5a-c) and other cysteine-proteases^43^. Since CTSF showed stronger inhibitory effects in cell culture, we decided to use AAV-hSyn-Cst7. Immunofluorescence confirmed neuronal CTSF expression (Fig. 5e), which resulted in reduced LGMN maturation and strongly reduced activity (Fig. 5f,g). Furthermore, as a consequence of the inhibition of LGMN activity, TDP-43 processing was also reduced to levels seen in wt co-culture (Fig. 5h).

To investigate potential shuttling of LGMN from microglia to neurons, we initially investigated whether LGMN can be secreted and found enhanced LGMN secretion in *GRN* ko hiMGL compared to wt hiMGL both co-cultured with wt hiNE (Fig. 5i). Conditioned media from these co-cultures were added to *GRN* wt hiNE monocultures (Fig. 5j). Transfer of conditioned media derived from co-cultured ko hiMGL and wt hiNE led to increased LGMN proteolytic activity in lysates of mono-cultured wt hiNE (Fig. 5j,k). This indicates that human neurons can internalize secreted LGMN further supporting our findings that secreted LGMN from microglia can lead to enhanced TDP- 43 processing in neurons.

**Fig. 5:**
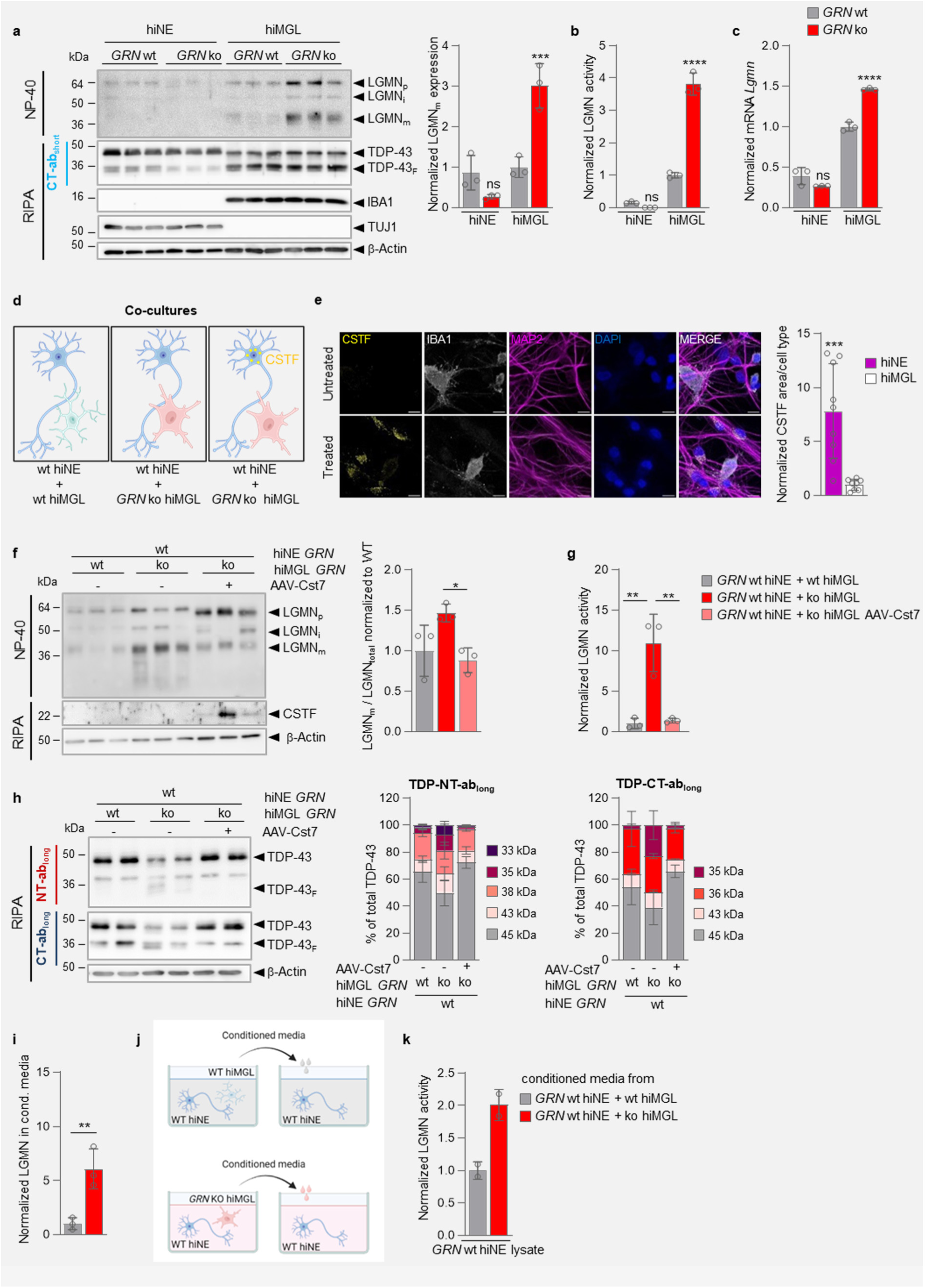
Elevated LGMN maturation and activity in human induced pluripotent stem cells (hiPSC)-derived microglia results in enhanced TDP-43 processing. **a,** Representative immunoblot for LGMN, TDP-43, IBA1, TUJ1, and β-actin of *GRN* wt and ko hiPSC-derived microglia (hiMG) or neurons (hiNE). LGMN_p_, LGMN_m_, full-length TDP-43, and TDP-43 fragments (TDP-43_F_) are indicated. Quantification of LGMN_m_ from the immunoblot shown as mean normalized to wt hiMGL ± s.d. (n=3). **b,** *In vitro* LGMN activity of lysates from monocultured wt and *GRN* ko hiMGL and hiNE; mean normalized to wt hiMGL ± s.d. (n=3). **c,** qPCR of *LGMN* mRNA expression in wt and *GRN*-deficient (*GRN* ko) hiMGL or hiNE normalized to wt hiMGL, mean ± s.d. (n=3). **D**, Scheme for **e-i**: 3-week co-culture of wt hiNE with *GRN* wt hiMGL, ko hiMGL, or ko hiMGL and AAV-Cst7 (cystatin F expressing (CSTF)) transduction. **e**, Immunofluorescence of CSTF (yellow), neurons (MAP2, magenta), and microglia (IBA1, white) with nuclear staining (DAPI, blue) upon AAV transduction (scale bar 15 µm). Quantification of CSTF area normalized over MAP2 area (hiNE) or IBA1 area (hiMGL). Data are mean ± s.d. of image replicates (n=9). **f**, Immunoblotting of LGMN, CSTF, and β-actin to verify equal loading. Quantification of LGMN_m_ over total LGMN; mean normalized to wt hiMGL co-culture ± s.d. (n=3). **g**, LGMN activity normalized to hiNE co-cultured with wt hiMGL shown as mean ± s.d. (n=3). **h**, Immunoblotting of TDP-43, using N-terminal (NT-ab_long_) or C-terminal (CT-ab_long_) anti-TDP-43 antibodies, and β-actin. Quantification of indicated TDP-43 fragments (TDP-43_F_) and TDP-43 holoprotein for both antibodies; mean normalized to wt hiMGL co-culture (n=2). **i**, ELISA of LGMN detected in the media; mean normalized to wt hiMGL co-culture ± s.d. (n=3). **j**, Scheme for (**k**); monocultured *GRN* wt hiNE treated for 24 h with conditioned media from either 3 weeks *GRN* wt hiNE and wt hiMGL co-cultures or *GRN* wt hiNE and ko hiMGL co-cultures. **k**, LGMN activity of treated monocultured *GRN* wt hiNE, normalized to *GRN* wt hiNE treated with conditioned media from wt co-cultures (n=2). Statistical significance was determined using one-way ANOVA and Tukey’s post hoc test: ns, not significant; \**P*<0.05, \*\**P*<0.01, \*\*\**P*<0.001, \*\*\*\**P*<0.0001. *P* values and statistical source data are provided.

To further validate the unique role of microglia-derived LGMN in TDP-43 processing, we treated the co-culture with a highly selective small molecule LGMN inhibitor (Extended Data Fig. 6). Upon three weeks of treatment, full-length TDP-43 was increased and TDP-43 fragments were significantly reduced. The remaining TDP-43 processing product of slightly higher molecular weight appears to be generated in a LGMN independent pathway (Fig. 6a,b). Pro-LGMN and mature LGMN were both strongly increased upon inhibitor treatment (Fig. 6a,c). Together with an approximately 90% reduction of the proteolytic activity (Fig. 6d), this suggests reduced self-degradation of LGMN. Taken together, these findings indicate that ko hiMGL release LGMN, which is taken up by neurons, where it then mediates pathological processing of TDP-43.

**Fig. 6:**
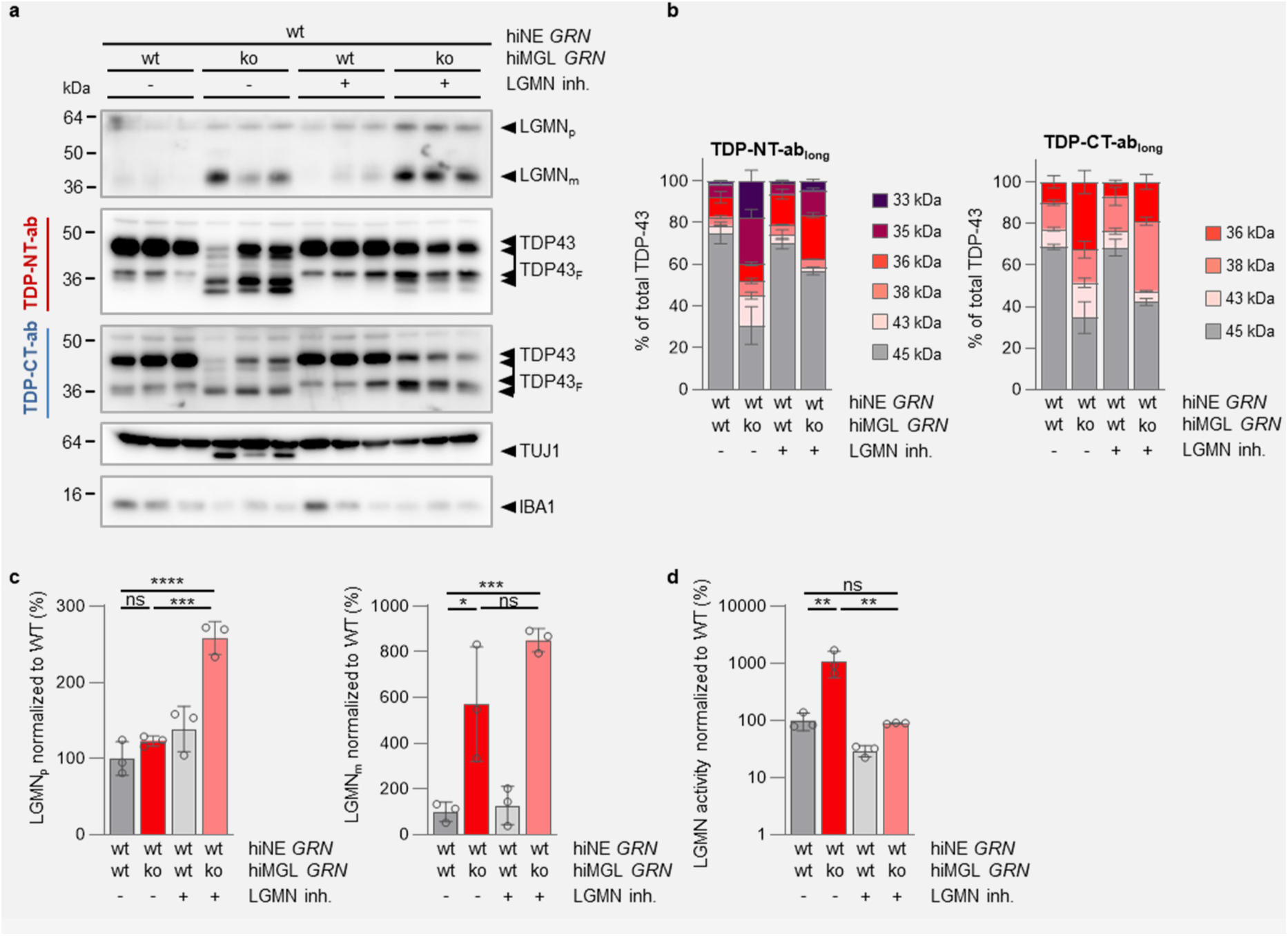
Inhibition of LGMN activity diminishes TDP-43 processing. **a**, Immunoblot of a 3 weeks co-culture of *GRN* wt hiNE with either *GRN* wt or ko hiMGL with or without treatment with a specific small-molecule LGMN inhibitor (inh.) every 48 h (n=3). LGMN (LGMNp and LGMNm), TDP-43 (full length and TDP-43F), TUJ1 and IBA1 were probed. **b**, Quantification of full-length TDP-43 and fragments in (**a**) shown as mean normalized to untreated *GRN* wt hiNE/ *GRN* wt hiMGL co-cultures ± s.d. (n=3). **c**, Quantification of LGMNp and LGMNm in (**a**) shown as mean normalized to untreated *GRN* wt hiNE/ *GRN* wt hiMGL cocultures ± s.d. (n=3). **d**, LGMN activity of the co-cultures in % (log scale) normalized to untreated *GRN* wt hiNE/ *GRN* wt hiMGL co-cultures ± s.d. (n=3). Statistical significance was determined using one-way ANOVA and Tukey’s post hoc test: ns, not significant; \**P*<0.05, \*\**P*<0.01, \*\*\**P*<0.001, \*\*\*\**P*<0.0001. *P* values and statistical source data are provided.

### LGMN overexpression in mice results in TDP-43 pathology and motor deficits

As inhibition of LGMN activity in neurons reduced pathological processing of TDP-43, we assumed that TDP-43 pathology could be conversely enhanced *in vivo* by overexpressing LGMN. To prove if increased hLGMN expression in the brain is sufficient to induce FTLD-related pathology, 6-7-month-old *Grn* ko mice were treated *via* tail vein injection with AAV-PhP.eB-CAG-h*LGMN*, a serotype that was shown to have high brain penetrance as well as transduction of neurons and astrocytes^44^. Three months post injection, we observed substantial overexpression of hLGMN (Fig. 7a) as well as a 13-fold increase of LGMN activity in total brain lysates (Fig. 7b). hLGMN was expressed in all brain areas (Extended Data Fig. 7a). hLGMN expression was predominantly found in neurons, and to a lesser extent in astrocytes and microglia (Extended Data Fig. 7b). In line with our *ex vivo* findings, upon hLGMN overexpression in *Grn* ko mice, we detected enhanced generation of TDP-43 fragments (Fig. 7c) and a strong accumulation of insoluble abnormally phosphorylated TDP-43 (pTDP-43) (Fig, 7c). Abnormal processing and pathological phosphorylation of TDP-43 was accompanied by motor deficits as shown by increased hind limb clasping (Fig. 7d) as well as progressing deficits in rotarod performance over time (Fig. 7e). These phenotypes resembled the phenotypes observed in the previously described *Tmem106b* ko/ *Grn* ko FTLD model^45^, suggesting that LGMN overexpression in *Grn* ko can induce an FTLD-like pathology. Finally, LGMN overexpression led to a 10-fold increase in plasma neurofilament light chain (NfL) (Fig. 7f), a biomarker for neuronal damage also used in clinical studies^46,47^. Taken together, these results suggest that LGMN is a main driver of TDP-43 pathology *in vivo*.

**Fig. 7:**
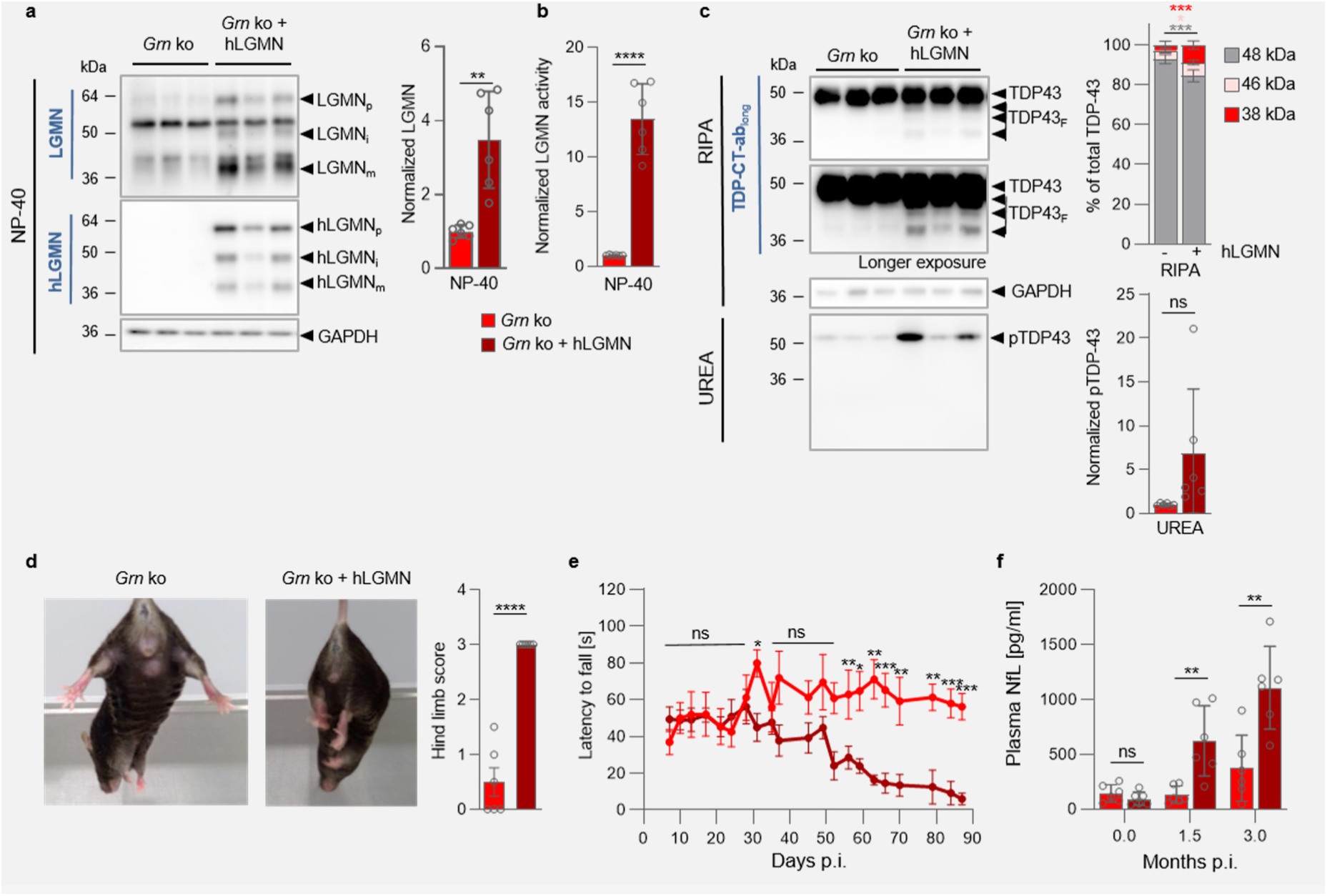
Overexpression of hLGMN in *Grn* ko mice leads to TDP-43 pathology, motor deficits and increased plasma NfL levels. **a**, Representative immunoblots for total LGMN and human LGMN (hLGMN) in total brain homogenate (NP40) of *Grn* KO mice, three months p.i. and age-matched non-injected controls, GAPDH verified equal loading. LGMN expression is quantified and normalized to *Grn* ko (n=3 mice per condition, mean ± s.d) **b**, *In vitro* LGMN activity assay of total brain homogenate of AAV-injected *Grn* ko mice and age-matched non-injected controls (n=6 mice per condition, mean ± s.d). **c**, Representative immunoblot of TDP-43 in the RIPA soluble fraction of total brain lysates and pTDP-43 in the urea-soluble fraction, GAPDH verified equal loading. For quantification of TDP-43 processing in the RIPA fraction, the signal of indicated TDP-43 fragments was normalized to the total TDP-43 signal (n=6 mice per condition, mean ± s.d., multiple t-tests with FDR correction). pTDP-43 accumulation in the urea fraction was quantified and normalized to *Grn* ko (n=6 mice per condition, mean ± s.d.), **d**, Representative images show assessment of hind-limb clasping phenotype in hLGMN overexpressing mice and age-matched non-injected *Grn* ko mice (n=6 mice per condition). The clasping phenotype was quantified (n=6 mice per condition, mean ± s.e.m.). **e**, Longitudinal rotarod performance of *Grn* ko mice overexpressing hLGMN and age-matched controls (n=6 for *Grn* ko + hLGMN, n=6 for *Grn* ko control mice, mean ± s.e.m., multiple t-tests with FDR correction). **f**, Plasma NfL levels were measured before injection,1.5 months p.i and terminally. (n=6, mean ± s.d.).

## Discussion

Our findings demonstrate that enhanced LGMN activity provides a direct link between *GRN* haploinsufficiency and TDP-43 pathology. In various model systems, including two independent PGRN-deficient mouse lines, MEF, primary mouse- and hiPSC-derived microglia, and finally FTLD-G*RN* patient brains, we consistently found elevated LGMN maturation and activity. Upon co-incubation of LGMN and PGRN *in vitro*, we observed that PGRN, but not granulin peptides, delays LGMN maturation in a dose-dependent manner. Thus, PGRN slows LGMN maturation, presumably in a chaperone-like manner during trafficking to lysosomes. The inhibitory effect is balanced by LGMN itself, as it hydrolyses PGRN into non-inhibitory granulin peptides. We found that granulin peptides are reduced in *Lgmn* ko mice, thus LGMN is involved in granulin peptide generation *in vivo*. This is further supported by recent *in vitro* data by Mohan et al., who investigated the role of various lysosomal proteases in the differential processing of PGRN^29^.

In addition, pathologically enhanced LGMN activity leads to increased processing of lysosomal cathepsins, which, consistent with previous findings^15,18,48^, may result in dysregulation and malfunction of microglial lysosomes. Considering the potential of several lysosomal cathepsins to process PGRN as well^27-29,49^, this could further accelerate the pathology. Based on our findings that both, knockdown of LGMN or selective inhibition of LGMN activity ameliorates pathological TDP-43 processing, whereas overexpression accelerates TDP-43 pathology, LGMN might be a main driver of FTLD-*GRN* pathology. Fragment formation is disease relevant, since proteolytically cleaved TDP-43 fragments are not only enriched in TDP-43 deposits, but such fragments are also known to be required for seeding and spreading of pathological TDP-43^50^. A contribution of other proteases activated by LGMN cannot be excluded though, as the overexpression of the more general protease inhibitor CSTF showed a stronger rescue than the selective small molecule LGMN inhibitor in our hiPSC co-culture system.

Strikingly, overexpression of LGMN in *Grn* ko mice caused a severe worsening of disease phenotypes, including processing, phosphorylation and aggregation of TDP-43. Similar effects of overexpressed LGMN have been proposed for tau processing and phosphorylation in Alzheimer’s disease^51,52^. Phosphorylation of tau might be affected by elevated LGMN levels which results in reduced protein phosphatase 2a (PP2A) stability and as a consequence in hyperphosphorylated tau^51^. Whether this mechanism plays a role for TDP-43 phosphorylation needs to be further investigated. In addition, in *Grn* ko mice, overexpressed LGMN resulted in increased plasma NfL levels and worsened rotarod performance. Elevated NfL levels in serum and CSF of FTLD patients correlate with disease severity and brain atrophy, with particular high levels in symptomatic FTLD-*GRN* patients^53,54^. TDP-43 pathology results in motor deficits in amyotrophic lateral sclerosis, but also FTLD and specifically FTLD-*GRN* patients can present with motor neuron disease-like syndromes^55^. Since TDP-43 cleavage removes the nuclear localization signal from the C-terminal fragments, it will further enhance cytoplasmic accumulation and aggregation of TDP-43. LGMN secretion has already been described in the periphery^36^. Here, LGMN is mainly expressed by immune cells and upregulated upon stress or in tumor microenvironment. In these conditions its pro-form was also found to be secreted^36^. The secretion of LGMN by activated microglia and further functional consequences need to be studied in more detail in the future. Together, our findings strongly suggest that the pro-LGMN is released by PGRN deficient microglia and internalized by neurons, where it is activated and capable of processing TDP-43, thus resolving the spatial paradox of the differential expression of LGMN and PGRN predominantly in microglia, and TDP-43 pathology primarily occurring in neurons^3^ (Fig. 8). In which cellular compartment LGMN gains access to TDP-43 remains to be discovered. To reach cytoplasmic TDP-43 mature LGMN might be released by damaged lysosomes^16^. Although active LGMN is unstable at the neutral pH of the cytosol, it might be stabilized by active center interacting proteins^36^. One could speculate that re-activation occurs by release of the binding partner in an acidic microenvironment. A more likely mechanism might be the direction of cytoplasmic TDP-43 towards autophagic degradation^56,57^ and cleavage by LGMN in autolysosomes. TDP-43 fragments might then escape damaged, leaky lysosomes^16^ and accumulate in the cytoplasm.

**Fig. 8:**
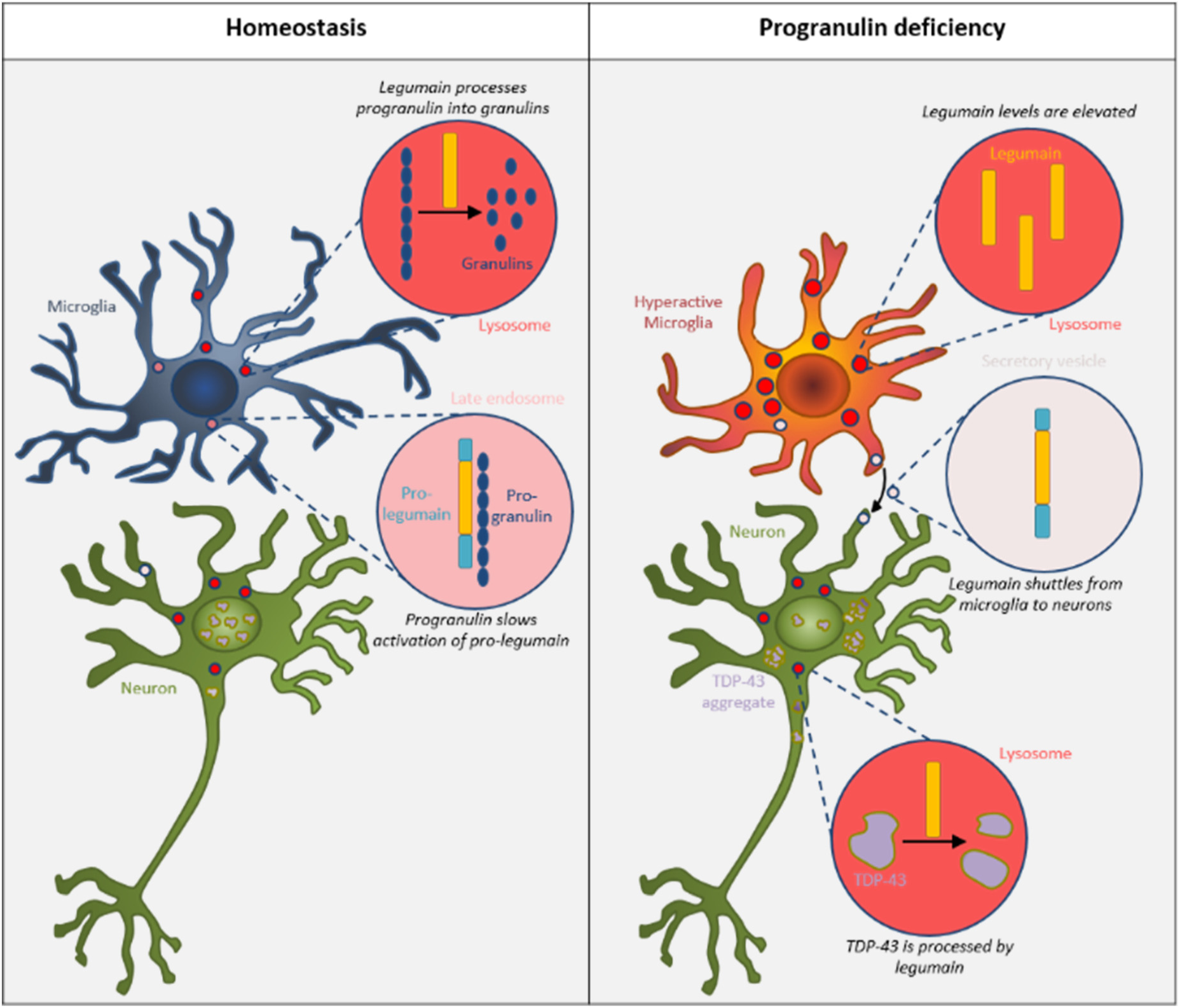
Increased legumain activity caused by progranulin deficiency drives TDP-43 pathology. Suggested pathomechanism: In homeostasis, PGRN slows the maturation of LGMN, which is regulated in a negative feed-back loop *via* processing of PGRN into single granulins by LGMN. Upon PGRN loss-of-function, lysosomal function is impaired and microglia become hyperactivated. In this context, LGMN is not regulated by PGRN anymore and its protein levels and activity increase. The pro-form of LGMN is secreted by activated microglia and taken up by neurons (mechanism unknown), where it hyperprocesses TDP-43 and thereby drives TDP-43 pathology.

PGRN-deficiency can cause microglial hyperactivation^10-12^ and triggers the complement cascade^58^ resulting in neuronal loss. How aberrantly elevated LGMN activity is linked to these processes remains to be investigated. As FTLD-*GRN* patients also present with co-pathology of other LGMN substrates, namely tau and α- synuclein^52,59,60^, it is conceivable that their accumulation and deposition may also be facilitated by aberrantly enhanced LGMN activity. In addition, LGMN activity was also found to be upregulated in the brains of patients with other neurodegenerative diseases, such as Alzheimer’s and Parkinson’s disease^52,61^. Monitoring LGMN activity in patient brains or the increase of LGMN specific cleavage products in the CSF could be useful biomarkers in the future to monitor disease progression and lysosomal dysfunction. Finally, since TDP-43 pathology and lysosomal dysfunction co-exist in other neurodegenerative diseases such as Alzheimer’s disease, Parkinson’s disease, limbic-predominant age-related TDP-43 encephalopathy (LATE), Lewy Body dementia and hippocampal sclerosis, our findings may reveal a disease overarching pathway, which could be modulated by LGMN inhibitors^62-66^.

## Methods

### Data reporting

No statistical methods were used to predetermine sample size. The investigators were not blinded to allocation during experiments and outcome assessment, except for animal motor function assessment tests.

### Ethical approval

All the work involving human tissues or mice was carried out in accordance with the Code of Ethics of the World Medical Association (Declaration of Helsinki). The use of human brain material was approved by the local ethic commission of the Ludwig-Maximilian’s-University, Munich. All animal experiments were performed in compliance with the national guidelines for animal protection in Germany and with the approval of the regional animal committee (Regierung von Oberbayern) directed by a veterinarian. Mice were kept in small groups under standard housing conditions at constant temperature of 21 ± 2°C, on a 12 h light / dark cycle, providing standard pellet food and water *ad libitum*. At Denali Therapeutics Inc. all mouse procedures adhered to regulations and protocols approved by Denali Therapeutics Institutional Animal Care and Use Committee. Mice were housed under a 12 h light / dark cycle and group housed when possible.

### Experimental mouse models

The *Grn* ko mouse line was provided by Dr. M. Nishihara (Department of Veterinary Physiology, The University of Tokyo)^67^, backcrossed to the C57BL/6J strain and published before^10,17,18,22,45^. Animal ages / sexes for each experiment are indicated in the source data tables.

### Overexpression of hLGMN in *Grn* ko mice

100 µl of AAV-PhP.eB-CAG-h*LGMN* with a titer of c=5*10^12^ vg/ml was applied *via* tail vein injection at an age of 6-7 months in male *Grn* ko mice. Non-injected age-matched male *Grn* ko mice were used as controls.

### Motor function assessment

#### Rotarod test

A rotarod (# 3376O4R, TSE Systems) was used to assess motor function. The speed was accelerated stepwise from 4-40 rpm within 300 s as previously described^45^. Mice were trained twice within one week before starting longitudinal rotarod measurements. The average latency to fall off the rotarod of three consecutive tests was calculated for each recording day.

#### Hind limb clasping test

Mice were recorded for 10 seconds while being lifted by their tail as previously described^63^. Mice were scored based on the degree of cramping of their hind limbs recorded on two different days at 3 months p.i.. Mice showing no cramping obtained a score of 0, cramping for less than 25% of recording time a score of 1, more than 25% of recording time a score of 2, or more than 50% of the recording time a score of 3.

### Mouse brain dissection and blood collection

Mice were sacrificed by CO2 inhalation or by deep/lethal anesthesia and perfused with ice cold PBS. Brain tissue dissected from adult mice was either snap frozen in liquid nitrogen, mechanically pulverized and stored at ™80°C for biochemical analysis, fixed in PFA for immunofluorescence staining or directly used for cell isolation.

Blood was collected 1 day before virus injection and at 1.5 months p.i. by facial vein puncture. Terminally, blood was collected before perfusion via cardiac puncture in EDTA-coated tubes and centrifuged (15,500 x g, 7 min, 4°C). For analysis the plasma fraction was used.

### Mouse brain cell isolation

Mice were sacrificed by CO2 inhalation or by deep/lethal anesthesia and perfused with ice cold PBS and directly subjected to microglia and astrocyte or neuron isolation. Neural cells were acutely isolated from adult mouse brain using MACS technology (Miltenyi Biotec). Brain tissue was dissociated into a single-cell suspension by enzymatic digestion using the Adult Brain Dissociation Kit P (# 130-107-677, Miltenyi Biotec) and the gentleMACS™ Octo Dissociator (# 614 130-096-427, Miltenyi Biotec) according to manufactureŕs instructions. Microglia and astrocyte isolation were performed from the same mouse brain. Briefly, the dissociated cell suspension was applied to a pre-wetted 100 μm cell strainer, cells were pelleted at 300 x g, 4°C, 10 min, pellets were washed twice and resuspended in 1 ml 0,5% (w/v) BSA/PBS. CD11b- positive microglia were magnetically labelled with 20 µl anti-CD11b MicroBeads (# 130- 049-601, Miltenyi Biotec) and incubated for 20 min in the dark at 4°C with gentle shaking. Cells were washed by adding 1−2 ml of BSA/PBS. The cell pellets were resuspended in 1 ml BSA/PBS and applied together with 1 ml BSA/PBS onto the prepared LS columns (# 130-042-401, Miltenyi Biotec) placed into a QuadroMACS™ Separator (# 130-091-051, Miltenyi Biotec). The columns were washed with 3 × 3 ml BSA/PBS. The flow-through and the first wash containing the unlabeled cells was collected for astrocyte isolation. The columns were removed from the magnetic field, and microglia were flushed out using 5 ml BSA/PBS. To obtain a higher purity, the microglia containing eluate was applied onto pre-wetted MS columns (# 130-042-201, Miltenyi Biotec) placed into an OctoMACS^TM^ Separator (# 130-042-108, Miltenyi Biotec), washed 3 x 0.5 ml and eluted with 1 ml BSA/PBS, then immediately stored on ice. For astrocyte isolation the microglia-depleted fraction including the first wash was pelleted and resuspended in 1 ml BSA/PBS, then 10 μl of FcR blocking reagent was added and incubated for 20 min, gentle shaking in the dark at 4°C, followed by adding

10 μl of anti-ACSA-2 MicroBeads (Anti-ACSA-2 MicroBead Kit, # 130-097-678, Miltenyi Biotec) and an additional incubation for 30 min. Cells were washed and astrocytes were magnetically separated as described for microglia. Neurons were isolated (Adult Neuron Isolation Kit mouse, # 130-126-603, Miltenyi Biotec) according to the manufacturer’s instructions including a red blood cell removal and a debris removal step (Adult Brain Dissociation Kit P, # 130-107-677, Miltenyi Biotec). Isolated cells were washed twice with D-PBS (# 14040-133, Thermo Fisher Scientific) to remove BSA, pellets were snap frozen in liquid nitrogen and stored at ™80°C until further biochemical analysis.

### Mouse embryonic fibroblasts (MEF), HeLa and HEK293T cell culture

All MEF were generated as described before^4^. MEF, HeLa, and HEK293T were cultured in Dulbecco’s modified Eagle’s medium (DMEM) with GlutaMAX^TM^-I (# 10566016, Thermo Fisher Scientific) supplemented with 10% (v/v) heat inactivated FBS (# F7524, Sigma-Aldrich), 100 U/ml penicillin, and 100 µg/ml streptomycin (# 15140148, Thermo Fisher Scientific), and if required 1% non-essential amino acids (# 11140050, Thermo Fisher Scientific).

### Primary culture of mouse microglia

Microglia were isolated from P11 pups using the MACS Technology (Miltenyi Biotec) as described above. Isolation was carried out under sterile conditions. 60,000 cells were plated in DMEM/F-12, HEPES (# 11594426, Thermo Fisher Scientific) supplemented with 10% (v/v) heat inactivated FBS (# F7524, Sigma-Aldrich), 100 U/ml penicillin, and 100 µg/ml streptomycin (# 15140148, Thermo Fisher Scientific).

### Primary culture of mouse hippocampal neurons

Mouse hippocampal neurons were generated as described previously^45^. In brief, E18 embryos were collected from CO2-euthanized C57BL/6J wildtype mice. The hippocampi were isolated and dissected in ice-cold dissection buffer (HBSS, 1 mM sodium pyruvate, 10 mM HEPES pH 7.2, all from Thermo Fisher Scientific). Single cell suspension was obtained by pelleting the tissue (300 x g, 3 min, RT) and incubating it for 15 min at 37°C with 0.125% trypsin (# 25200-072, Thermo Fisher Scientific) and 25 U/ml benzonase (# E1014-25KU, Sigma-Aldrich) in DMEM with GlutaMAX^TM^-I. After centrifugation (300 x g, 3 min, RT), tissue was washed in DMEM with GlutaMAX^TM^-I and pelleted again (300 x g, 3 min, RT) before trituration in culture media (neurobasal media, # 21103-049, Thermo Fisher Scientific) with 100 U/ml penicillin, 100 µg/ml streptomycin, 0.5 mM L-glutamine (# 25030-024, Thermo Fisher Scientific) and 1x B-27 supplement (# 17504-044, Thermo Fisher Scientific) with 25 U/ml benzonase and subsequent centrifugation (300 x g, 3 min, RT). The single cells were plated onto poly-D-lysine-(10 µg/ml, # P7280-5MG, Sigma-Aldrich) and laminin-(10 µg/ml, # L2020-1MG, Sigma-Aldrich) coated plates in culture media. On DIV 7, 20% media was added and on DIV 14, cells were used for experiments.

### Gene expression analysis

For quantitative real time PCR (qRT-PCR) approximately 10-20 mg of powdered mouse brain homogenates or between 400,000 and 2 million hiPSC-derived MGL or hNE were subjected to total RNA preparation using the QIAshredder and RNeasy Mini Kit (# 79656, Qiagen) according to manufacturer’s instructions. 0.1-1 µg of RNA was reverse transcribed into cDNA using M-MLV reverse transcriptase (Promega) and oligo(dT) primers (Life Technologies). The following primer sets from Integrated DNA Technologies were used: mouse *Lgmn* Mm.PT.58.5122210 (Exon boundary 10 to 11), mouse *Gapdh* Mm.PT.39a.1 (Exon boundary 2 to 3), human *LGMN* Hs.PT.58.21235685 (Exon boundary 3 to 4) and human *RPL22* Hs.PT.58.41088691(Exon boundary 3 to 4).

For quantification, cDNA levels were normalized to mouse *Gapdh* or human *RPL22* cDNA and relative transcription levels were analyzed using the comparative ΔΔCt method (7500 Software V2.0.5, Applied Biosystems, Life Technologies). The *Lgmn* mRNA level of the microglia-enriched fraction was obtained from a NanoString dataset (Omnibus GSE129709 https://www.ncbi.nlm.nih.gov/geo/query/acc.cgi?acc=GSE129709)^10^.

### Cloning of m*Lgmn*

The mouse *Lgmn* cDNA was amplified from MEF cDNA prepared as described for “gene expression analysis”, following primers were used: *Lgmn* (XhoI) forward 5’- CCCGCTCGAGGCCACCATGACCTGGAGAGTGG-3’ *Lgmn* (EcoRV) reverse 5’-AGCTTTGATATCTCAGTAGTGACTAAGAC-3’. The amplified *Lgmn* cDNA was sub-cloned into the XhoI/EcoRV sites of pcDNA3.1/Zeo(-). The *Lgmn* mutation leading to an inactive LGMN C191A variant^40,68^ was introduced by site-directed mutagenesis using the QuickChange Site-Directed Mutagenesis Kit (# 200519, Agilent Technologies) according to manufacturer’s instruction. The Cys coding base triplet TGT was exchanged to GCT coding for Ala using following primers: *Lgmn*(mt)F 5’- GGTGTTCTACATTGAAGCTGCTGAGTCTGGCTCCATGATGAACC-3’-*Lgmn*(mt)R 5’-GGTTCATCATGGAGCCAGACTCAGCAGCTTCAATGTAGAACACC-3’.

### Design and production of adeno-associated virus

*AAV-hSyn-Cst7.* A 603 bp long nucleotide encoding for cystatin-7 was cloned into the BamHI/EcoRI sites of pAAV-hSyn-EGFP (# 50465, Addgene) to receive the plasmid pAAV-hSyn-*Cst7*. To generate double-stranded adeno-associated virus of serotype 9 (AAV9), the plasmid and helper plasmid were transfected into HEK293T cells using polyethylenimine (# 24765, Polysciences). The virus was harvested after 72 h and purified from benzonase-treated cell lysates via an iodixanol density gradient (OptiPrep, Fresenius Kabi Norge). After rebuffering in lactose AAV titers were determined by real-time PCS on vector genomes using the SYBR Green Master Mix (Roche Molecular Systems).

*AAV-PhP.eB-CAG-hLGMN.* The AAV-PhP.eB-CAG-*hLGMN* vector used in this study was produced under research-grade conditions at SignaGen by triple transfection of adherent HEK293T cells. Briefly, HEK293T were transiently transfected after 48 h of culture using the corresponding Gene of Interest (GOI) plasmid, the corresponding capsid plasmid (AAV-PHP.eB), and the helper plasmid and PolyJet DNA transfection reagent (SignaGen) as transfection reagent. 72 h after transfection, cells were detached and harvested using sterile PBS. Harvested cells were lysed by three freeze/thaw cycles before an overnight benzonase treatment at 37 °C to digest residual DNA followed by high salt treatment for 1 h at 37°C. The crude lysate was further clarified by low-speed centrifugation. Full capsid AAVs were purified by ultracentrifugation in a cesium chloride (CsCl) gradient. A diafiltration step was then performed to concentrate the purified AAVs and reformulate them into PBS + 0.005% poloxamer 188 (# 13-901-CI, Corning) using Amicon Ultra-4 filter units (# UFC8100, Merck) with a 100 kDa MWCO membrane. Finally, a sterile filtration through a 0.22 μm membrane filter was performed. Vector genome titer was determined by qPCR using an ITR-specific primer-probe set. The final purified products were characterized with assays to measure identity, potency, and purity.

### Lipofectamine 2000 mediated cDNA transfection

For transfection of HeLa and HEK293T cells Lipofectamine 2000 (# 11668030, Thermo Fisher Scientific) was used according to the manufacture’s instruction. Briefly, 500 µl of 4 µg/ml plasmid solution in media was mixed with 500 µl DMEM with GlutaMAX^TM^-I containing 20 µl Lipofectamine 2000 and incubated for 20 min at RT before dropwise addition onto HeLa cells at 50-70% confluency. After 6 h, the media was exchanged and after 36 h the media was replaced with 4 ml Opti-MEM^®^ with GlutaMAX^TM^-I media (# 51985-026, Thermo Fisher Scientific). Media and cells were collected 48 h after transfection.

### siRNA Transfection

*Lgmn* siRNA (# M-044015-01-0005, siGENOME SMART pool, Dharmacon), control siRNA (#D-001210-04-20, siGenome non-targetting pool 4, Dharmacon) and mock reverse transfections were carried out with Lipofectamine^TM^ RNAiMAX (# 13778030, Thermo Fisher Scientific). For each 10 cm dish 3 µl siRNA (20 µM), were diluted in 600 µl Opti-MEM^TM^ (# 31985-062, Thermo Fisher Scientific) before adding 30 µl Lipofectamine^TM^ RNAiMAX and then for complex formation incubated for 15 min at RT.

Trypsinized and washed MEF cells were added to reach a final volume of 6 ml culture media without antibiotics and a final siRNA concentration of 10 nM. The next day, media were exchanged to fresh culture media and 96 h after transfection cells were harvested and pellets were washed in PBS, snap frozen in liquid nitrogen and stored at ™80°C.

### Human iPSC culture

HiPSC experiments were performed in accordance with relevant guidelines and regulations. The female iPSC line A18944 was purchased from Thermo Fisher (# A18945), grown in Essential 8 Flex Medium (# A2858501, Thermo Fisher Scientific) on VTN-coated (# A14700, Thermo Fisher Scientific) cell culture plates with 5% CO2 at 37°C, and split as small clumps twice a week after a 5 min incubation in PBS/EDTA. The *GRN* ko iPSC line was described before^22^.

### Differentiation of human iPSC-derived to cortical hiMGL, hiNE, co-culture of hiNE with hiMGL and treatment with AAV-Cst7 or LGMN inhibitor

HiPSC-derived hiMGL were differentiated as previously described^22^ following the protocol by Abud et al.^69^ with modifications to improve yield and efficiency. HiMGL were used for monoculture experiments on day 16 of the differentiation. HiPSC-derived hiNE were differentiated as previously published^70^ and used following day 60 of differentiation. HiNE were transduced with AAV-Cst7 at 500,000 gc/cell one week before co-culture with microglia. For co-culture experiments, hiMGL were added to differentiated neurons on day 12 of hiMGL differentiation and co-cultured for three weeks before collection for analysis. For LGMN inhibitor treatments, co-cultures were treated every 48 h from the start of co-culture with either 1 µM LGMN inhibitor (Roche) or a DMSO control for three weeks.

### Conditioned media treatment of monocultured hiNE

Monocultured hiNE were maintained along with co-cultures, as described above. Half of the media was taken out of the monocultured hiNE and media from the co-cultures was added before collecting the monocultured hiNE 24 h later.

### LGMN *in vitro* activity assay

A fluorescence-based activity assay was used to assess LGMN proteolytic activity of recombinant LGMN, or LGMN activity in lysates and media of the respective cells. Cell pellets or aliquots of powdered brain tissues were homogenized in LGMN-lysis buffer (50 mM sodium citrate pH 5.0, 0.8% NP-40, 1 mM DTT), incubated 15 min (cell lysates) or 20 min (brain lysates) on ice, followed by a 15 min centrifugation at 15,000 x g, 4°C. Protein concentration was determined using the BCA protein assay (Pierce, Thermo Scientific) and indicated amounts of protein in 50 µl LGMN-lysis buffer were pre-incubated in black 96-well plates (FluoroNunc) at 37°C for 10 min, then 50 µl LGMN assay buffer (50 mM sodium citrate pH 5.0, 1 mM EDTA pH 8.0, 1 mM DTT and 200 µM substrate (Z-Ala-Ala-Asn-AMC (# l-1865.0050, BACHEM))) prewarmed at 37°C was added. Cleavage of the quenched fluorescence substrate was continuously measured for 30 min following the increase of the fluorescence signal (excitation, 390 nm; emission, 460 nm) using the Fluoroskan Ascent FL plate reader (Labsystems). The relative enzyme activity was calculated for a period of time with linear substrate turnover.

### *In vitro* activation assays of recombinant pro-LGMN (rLGMN) or of the secreted pro-LGMN

**(sLGMN)** Prior to LGMN activity assays mouse rLGMN (# 2058-CY-010, R&D Systems) or sLGMN proform needs to be autocatalytically activated at acidic pH. 50 ng rLGMN or 25 µl of the collected media of transfected HeLa cells or hiMGL/hiNE co-cultures was preincubated in 10 µl or 25 µl acidic activation buffer (100 mM sodium citrate pH 4.0 or 3.5, 1 mM EDTA, 50 mM NaCl) for 4 h or as indicated at 37°C. For the activity assay 40 µl LGMN-lysis buffer was added and LGMN activity was measured as described above. To investigate the inhibitory effect of PGRN on LGMN activity, 200 ng or the indicated amount of human recombinant PGRN (rPGRN, # 10826-H08H, Sino Biological Inc.) was either added to the activation assay or to the activity assay.

### hLGMN ELISA

hLGMN was measured by an ELISA protocol previously established for PGRN by our group using the MSD Platform^71^. The ELISA consists of streptavidin-coated 96-well plates (streptavidin Gold Plates, # L15SA, MSD), a biotinylated polyclonal goat anti-human LGMN capture antibody (0.5 μg/ml, # BAF2199, R&D Systems), a polyclonal rabbit anti-human LGMN detection antibody (0.25 µg/ml, # NBP1-87793, Novusbio) and a SULFO-TAG-labelled goat polyclonal anti-rabbit IgG secondary antibody (0.5 μg/ml, # R32AB, MSD). All antibodies were diluted in 0.5% bovine serum albumin (BSA) and 0.05% Tween 20 in PBS (pH = 7.4) buffer. Recombinant human legumain/asparginyl endopeptidase (# 2199-CY-010, R&D Systems) was used as a standard (18.31 to 2343.75 pg/ml). In brief, streptavidin-coated 96-well plates were blocked overnight at 4°C in blocking buffer, then capture antibody (25 µl per well) was incubated for 1 h at RT, followed by four times washing (0.05% Tween 20 in PBS) and incubation for 2 h at RT with the samples diluted 1:2.5 in assay buffer (0.25% BSA and 0.05% Tween 20 in PBS (pH7.4)) or the recombinant human LGMN standard dilution. Plates were again washed four times with washing buffer before incubation for 1 hour at RT with the detector antibody. After four additional washing steps, plates were incubated with the secondary antibody for 1 h in the dark. Last, plates were washed four times with wash buffer followed by two washing steps in PBS. The electrochemical signal was developed by adding MSD Read buffer T (# R-92TC) and the light emission measured using the MSD MESO QuickPlex SQ120. The ELISA showed accurate results in spike recovery experiments (98%) and minimal measurement variation (CV = 4%) between replicates. For calculation of the sample concentration the MSD software Workbench v4 was used.

### Plasma Neurofilament light-chain (NfL) analysis

Plasma Nfl levels were measured with the Quanterix Simoa® NF-light^TM^ Reagent Kit (# 103186, Quanterix) following the manufacturer’s instructions as previously described (Reifschneider, 2022). Plasma NfL levels were run on a Simoa HDX instrument (Quanterix) and interpolated against a calibration curve included in the Quanterix assay kit.

### Generation of granulin peptides

Elastase generated granulin peptides were obtained by incubation of rPGRN with human neutrophil elastase (# RP-77526, Invitrogen). 50 ng/µl PGRN and 80 ng/µl elastase were incubated in 100 mM sodium citrate pH 6.0 overnight at 37°C, followed by buffer exchange to LGMN activation buffer. Aliquots were stored at ™80°C.

### Antibodies

| Antibodies | Dilution |  | Source |
| --- | --- | --- | --- |
|  | IB | IF |  |
| Mouse anti-β-actin | 1:5,000 | -- | # A 5316, Sigma-Aldrich |
| Rabbit anti-calnexin | 1:5,000 | -- | # SPA-860, Stressgene |
| Rabbit anti-cystatin-7 | 1:2,500 | 1:250 | # 50236, SinoBiological |
| Mouse anti-GAPDH | 1:10,000 | -- | # AM4300, Invitrogen |
| Goat anti-human IBA1 | -- | 1:250 | # ab5076, Abcam |
| Rabbit anti-human IBA1 | 1:500 | -- | # PA5-27436, Thermo Fisher Scientific |
| Rat anti-mouse LAMP1 | -- | 1:500 | # sc-19992, Santa Cruz |
| Sheep anti-mouse LGMN | 1:2,500 | 1:250 | # AF2058, R&D Systems |
| Goat anti-human LGMN | 1:1,000 | 1:250 | # AF2199, R&D Systems |
| Chicken anti- MAP2 | -- | 1:1,000 | # NB300-213, Novus Biologicals |
| Rat anti-mouse PGRN, (clone 8H10) | 1:50 | -- | generated<sup>72</sup> |
| Rabbit anti-human PGRN | 1:1,000 | -- | # 40-3400, Thermo Fisher Scientific |
| Rabbit anti-human PGRN / granulin-peptides | 1:1,000 | -- | # HPA008763, Atlas Antibodies |
| Rabbit anti-TDP-NTlong | 1:4,000 | 1:600 | # 10782-2-AP, Proteintech |
| Rabbit anti-TDP-CTlong | 1:1,000 | -- | # 12892-1-AP, Proteintech |
| Rabbit anti-TDP-CTshort | 1:1,000 | -- | # TIP-TD-P09, Cosmo Bio |
| Rabbit anti-pTDP-43 | 1:5,000 | -- | # 80007-1-RR, Proteintech |
| Mouse anti-tubulin $\alpha$ | 1:500 | -- | # T5168, Sigma-Aldrich |
| Mouse anti-tubulin β3 (TUJ) | 1:1,000 | -- | # 801201, BioLegend |

The following secondary antibodies were used for immunoblots: horseradish peroxidase-conjugated donkey anti-goat IgG (H+L) (Dianova, 1:5,000), donkey anti-sheep IgG goat (Jackson Immuno Research, 1:10,000), goat anti-mouse IgG (Promega, 1:10,000), goat anti-rabbit IgG (Promega, 1:20,000), goat anti-rat IgG + IgM (L+M) (Dianova, 1:5,000), and generated mouse anti-rat IgG2c (1:1,000).

### Immunohistochemistry

Cells seeded on coverslips in 24-well dish were fixed for 15 min in 4% paraformaldehyde and washed with PBS, then permeabilized for 10 min at room temperature in (0.1% Triton-X-100 in PBS for MEF and iPSCs, 0.1% Triton-X-100 and 0.1% saponin in PBS for *Cst7*-transduced hiN) and then washed with PBS again. SuperBlock (# 37515, Thermo Fisher Scientific) was used to block the coverslips for 1-2 h at RT. Primary antibodies were diluted in SuperBlock added overnight (iPSC) or for 2 h at 37°C (MEF). The coverslips were washed again in PBS and incubated in secondary antibodies (donkey anti-chicken Alexa 647 1:500, donkey anit-rat Alexa 647 1:500, donkey anti-sheep Alexa 555 1:500, donkey anti-rabbit Alexa 488 1:500, DAPI 1:5,000) diluted in SuperBlock for 90 min at RT in the dark. They were washed in PBS again and mounted with Fluoromount G (# 00-4958-02, Thermo Fisher Scientific).

### Protein analysis and immunoblotting

Snap frozen cell pellets or powdered brain homogenate were lysed in NP-40 LGMN- lysis buffer (50 mM sodium citrate pH 5.0, 0.8% NP-40, 1 mM DTT), NP-40 STEN-lysis buffer (150 mM NaCl, 50 mM Tris–HCl pH 7.6, 2.5 mM ETDA, 1 % NP40) or RIPA-lysis buffer (150 mM NaCl, 20 mM Tris–HCl pH 7.6, 2.5 mM ETDA, 1% NP-40, 0.1% sodium dodecyl sulfate (SDS), 0.5% sodium-desoxycholate) as indicated. NP-40 STEN- and RIPA-lysis buffer were supplemented with protease inhibitor cocktail (# P8340, Sigma-Aldrich) and phosphatase inhibitor (# 4906845001 PhosStop™, Sigma-Aldrich). Lysates were centrifuged for 20 min, 17,000 x g, 4°C. The protein concentration of the soluble fraction was determined using the BCA protein assay (Pierce, Thermo Fisher Scientific) and equal amount of protein were separated by SDS-PAGE and transferred to polyvinylidene difluoride membranes (Immobilon-P, Merck Millipore). Membranes were blocked for one hour in I-Block^TM^ (# T2015, Thermo Fisher Scientific). Proteins of interest were detected by the indicated primary antibodies followed by horseradish peroxidase-conjugated secondary antibodies detected by ECL Plus (# 32132×3, Pierce^TM^ ECL Plus Western Blotting Substrate, Thermo Fisher Scientific). For the quantitatively analysis, images were taken by a Luminescent Image Analyzer LAS-4000 (Fujifilm Life Science, Tokyo, Japan) and evaluated with the Multi GaugeV3.0 software (Fujifilm Life Science, Tokyo, Japan).

### Statistical analysis

Data are presented as mean ± s.d. or mean ± s.e.m. Statistical significance was calculated for comparison of two sample groups by unpaired, two-tailed Student’s t-test, and for multiple comparison by one-way ANOVA with Tukey’s or Dunnett’s post hoc test and indicated as ns, not significant; *P*>0.05; \**P*<0.05; \*\**P*<0.01; \*\*\**P*<0.001; \*\*\*\**P*< 0.0001. All data were analyzed using GraphPad Prism 7 (GraphPad Software Inc.).

## Data availability

All data and information are included in the manuscript. Supplementary information and source data are provided with this article. Data are available from corresponding authors upon reasonable request.

## Acknowledgement

We thank Ignacio Paris for image analysis, Lis de Weerd for help with tissue collection and the BELNEU consortium for clinical and pathological characterization for inclusion in the cohort of FTD patients, and the Institute Born-Bunge for the brain material. We thank Benedikt Wefers and Wolfgang Wurst for generating *Lgmn* ko mice. We further thank Joseph W. Lewcock for helpful discussions, Sarah De Vos and Kirk Henne for their inputs on study design, Ray Low for generation of PTV:PGRN material and Tim Earr for *in vivo* study assistance. Figures 4a and 5d&j were created with BioRender.com. This work was supported by a grant of the Alzheimer’s Association (to AC, CH, DP, MD) and the Deutsche Forschungsgemeinschaft (DFG, German Research Foundation) under Germany’s Excellence Strategy within the framework of the Munich Cluster for Systems Neurology (EXC 2145 SyNergy – ID 390857198). Parts of the project were also supported by the Koselleck Project (HA1737/16-1) of the DFG and the NOMIS foundation. CVB was supported by the Flemish Government initiated Methusalem excellence program, the Flanders Impulse Program on Networks for Dementia Research (VIND), the Research Foundation Flanders (FWO) and the Belgium Alzheimer Research Foundation (SAO). EW received a postdoctoral fellowship of the FWO.

## Author information

### Authors and Affiliations

Present address: Sophie Robinson, Cure Ventures, Boston, MA, USA. Todd Logan, BioMarin Pharmaceutical Inc., San Rafael, CA, USA. Anika Reifschneider, Bristol Myers Squibb, Munich, Germany.

## Author contributions

AC and CH conceived and designed the study and wrote the manuscript with the help of MTM, MR and SR. SR, MR and DP designed, performed and analyzed iPSC experiments. MR, DE, HR and AR performed and analyzed primary neuronal and microglial experiments. KB cloned LGMN constructs. MR performed and analyzed HeLa conditioned media-treated neuron experiments. KB and GW performed experiments with MEF, mouse brain and FTLD-*GRN* patient material and recombinant LGMN. MTM and MR performed *in vivo* mouse experiments and MTM analyzed the AAV-overexpression study in *Grn* ko mice. MJS, TL and GDP generated, MR analyzed the PTV:PGRN study. QM conducted the cystatin experiments. AA and SE generated and purified the AAV9-Cst7. CVB and EW provided validated FTLD-*GRN* patients brain samples. MD co-initiated the study and performed liver lysosome isolation on mice provided by AC, MTM and TR. SG and BB discovered and characterized the legumain inhibitor (LGMN inh). All authors commented on the manuscript.

These authors contributed equally:

Sophie Robinson, Marvin Reich and Maria-Teresa Mühlhofer

### Corresponding authors

Correspondence to Anja Capell or Christian Haass

## Ethics declarations

CH collaborates with Denali Therapeutics, participated on one advisory board meeting of Biogen and is a member of the advisory board of AviadoBio. DP is a scientific advisor of ISAR Bioscience. MJS, TL and GDP are full time employees and/or shareholders of Denali Therapeutics Inc. SG and BB are full time employees and/or shareholders of F. Hoffmann - La Roche Ltd.

## Additional Information

### Extended data figures and tables

**Extended Data Fig. 1:**
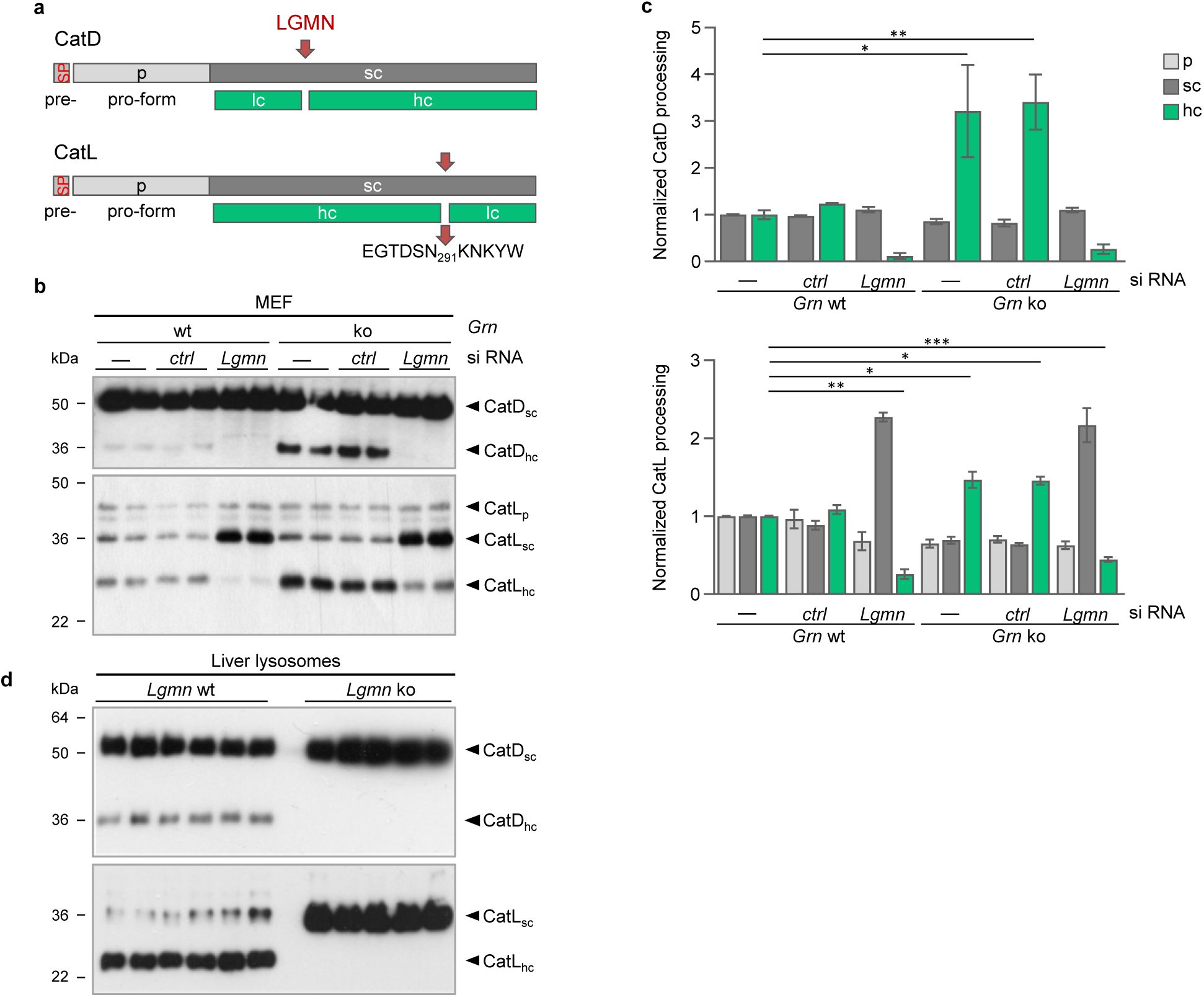
Elevated LGMN maturation and activity upon PGRN deficiency results in altered cathepsin processing. **a**, Processing of cathepsin D (CatD) and L (CatL) by LGMN is shown schematically. **b**, Representative immunoblots of MEF isolated from wt and *Grn* ko mice probed for CatD and CatL. MEF were either non-(-), control (*ctrl*) or *Lgmn* siRNA-transfected. CatD, and CatL, proform (p), single chain (sc), heavy chain (hc) are indicated. **c,** CatD and CatL maturation are quantified and normalized to wt (n=4). **d**, Representative immunoblots of liver lysosomes isolated from 6 months old *Lgmn* ko and wt mice probed for CatD and CatL. Data are mean ± s.d; unpaired two-tailed ttest, *p<0.05, **p<0.01, ****p<0.0001.

**Extended Data Fig. 2:**
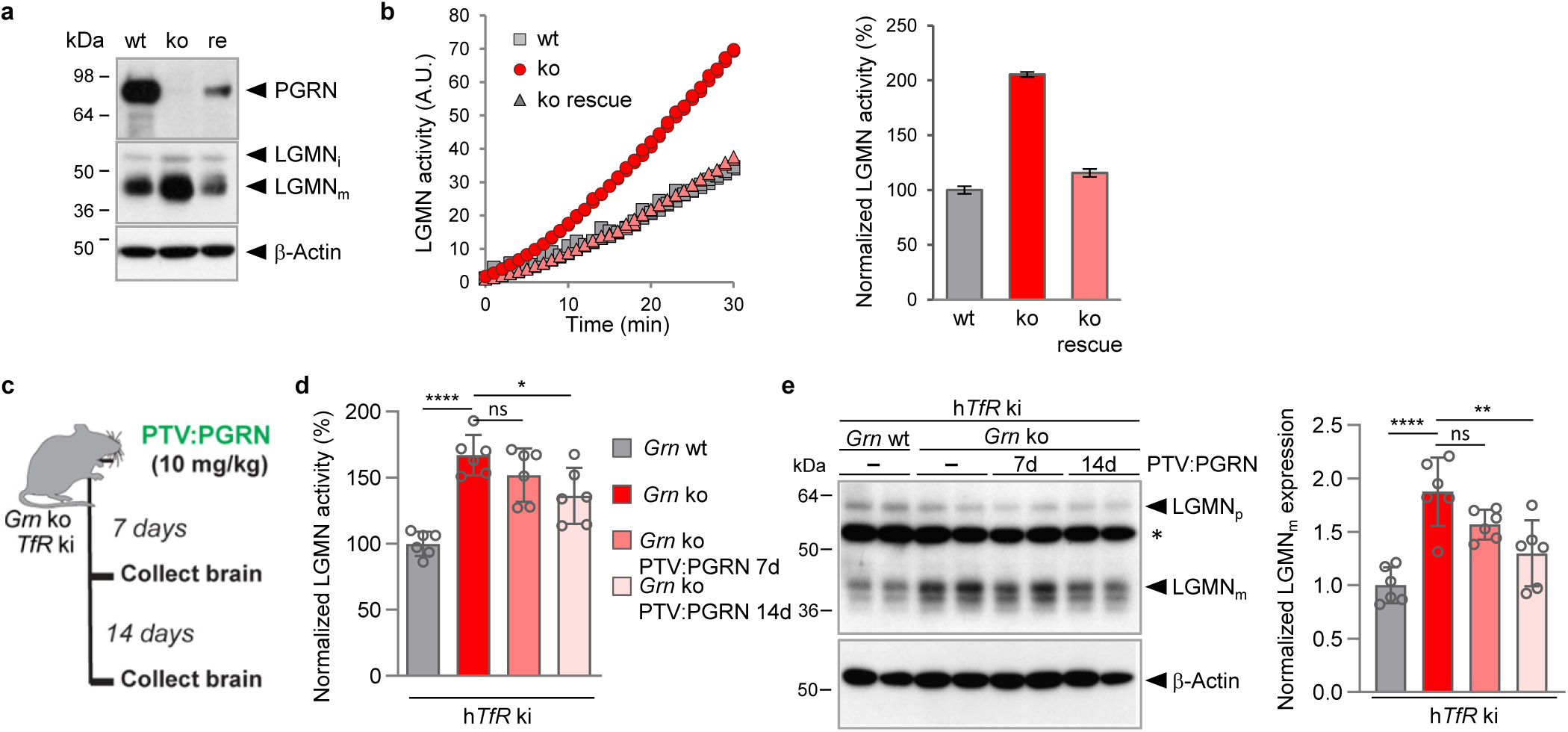
Enhanced LGMN activity of *Grn* ko mice and MEF is rescued upon PGRN expression. **a**, Representative immunoblots of MEF wt, *Grn* ko and *Grn* ko stably transfected with a mouse *Grn* cDNA probed for PGRN, LGMN, and β-actin to verify equal loading. **b**, LGMN substrate turnover and quantification of LGMN activity normalized to MEF wt of biologically independent duplicates is shown as mean ± s.d.. **c,** Schematic of PTV:PGRN single dosing study in *Grn* ko/h*TfR* ki. **d**, LGMN activity of a *Grn* wt, *Grn* ko, and *Grn* ko treated with a single 10 mg/kg i.v. dose of PTV:PGRN for 7 d and 14 d. Data indicate the mean normalized to wt. (n=6 mice per genotype and treatment, 3-4-month-old). **e**, Representative immunoblot for LGMN in total brain homogenate of wt and *Grn* ko non-treated and treated mice. Proform (LGMNp), mature form (LGMNm), unspecific band (asterisk) are indicated, β-actin verified equal loading. Quantification of LGMN immunoblot signals normalized to wt (n=6 mice per genotype and treatment). Statistical significance was determined using one-way ANOVA and Tukey’s post hoc test: ns, not significant; \**P*<0.05, \*\**P*<0.01, \*\*\*\**P*<0.0001. *P* values and statistical source data are provided.

**Extended Data Fig. 3:**
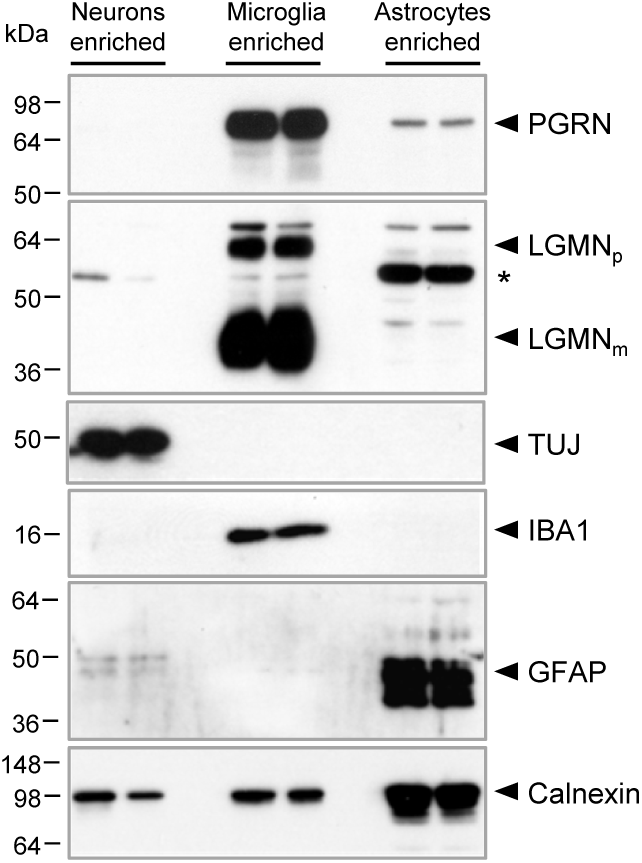
PGRN and LGMN are enriched in microglia. Representative immunoblots for PGRN and LGMN in microglia-, astrocyte-and neuron-enriched brain cell fractions isolated from 10-month-old mice. Enrichment of each cell fraction is verified by immunoblotting for IBA1, GFAP and TUJ. Equal amounts of protein were loaded for all cell types, calnexin expression varies between cell types. n=2 mice per cell type and genotype.

**Extended Data Fig. 4:**
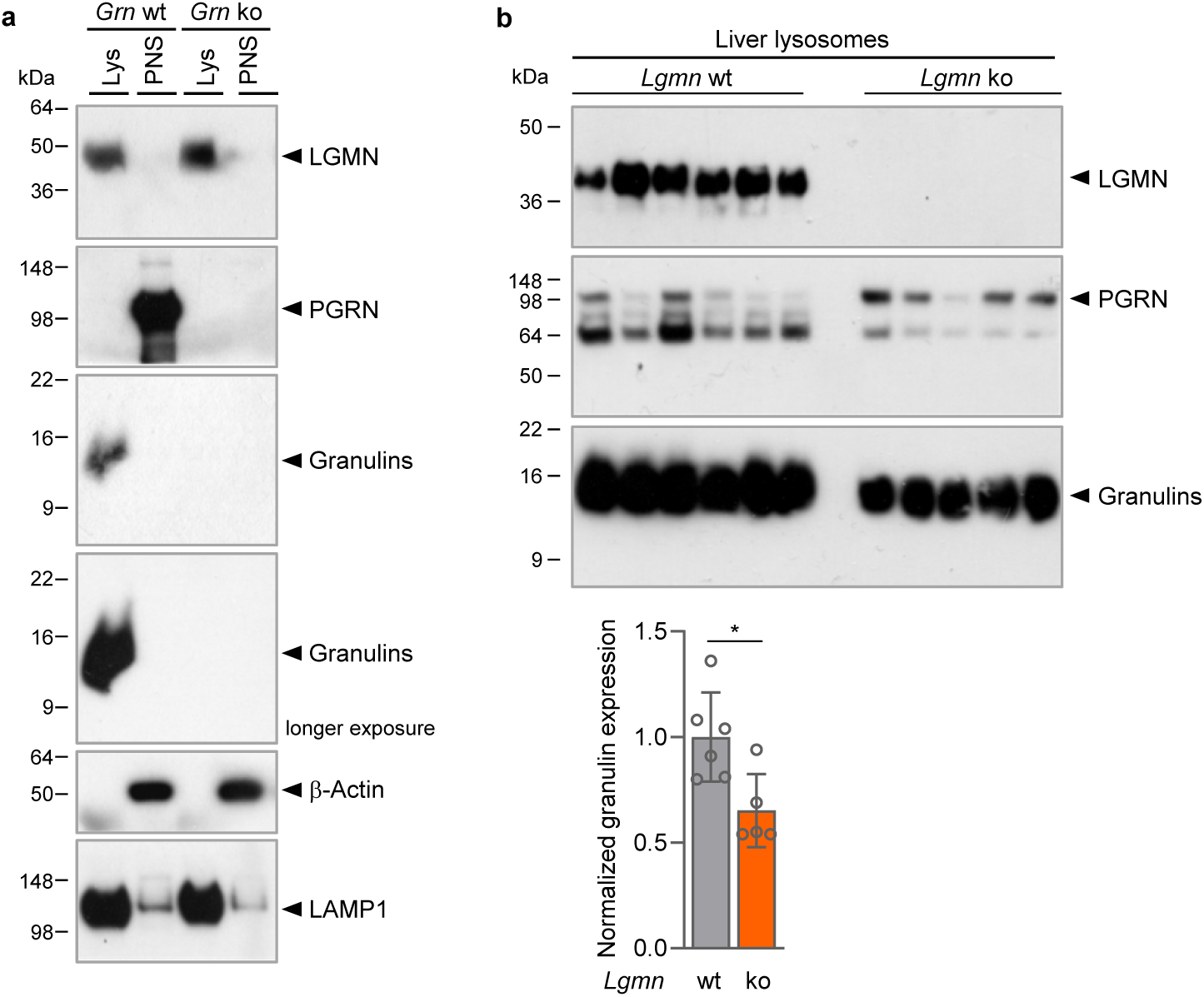
LGMN deficiency results in reduced generation of single granulins in liver lysosomes. **a**, Representative immunoblot of liver lysosome preparation of wt and *Grn* ko mice probed for LGMN, PGRN, granulins, with β-actin and LAMP1 as loading control for post nuclear supernatant (PNS) and lysosomes (Lys). **b**, Immunoblot of liver lysosome preparation of Lgmn wt and ko mice, probed for LGMN, PGRN and granulins. Data are mean ± s.d. of granulins in lysosome preparation from individual mice (n=5-6); unpaired two-tailed t-test; *p<0.05, *P* values and statistical source data are provided.

**Extended Data Fig. 5:**
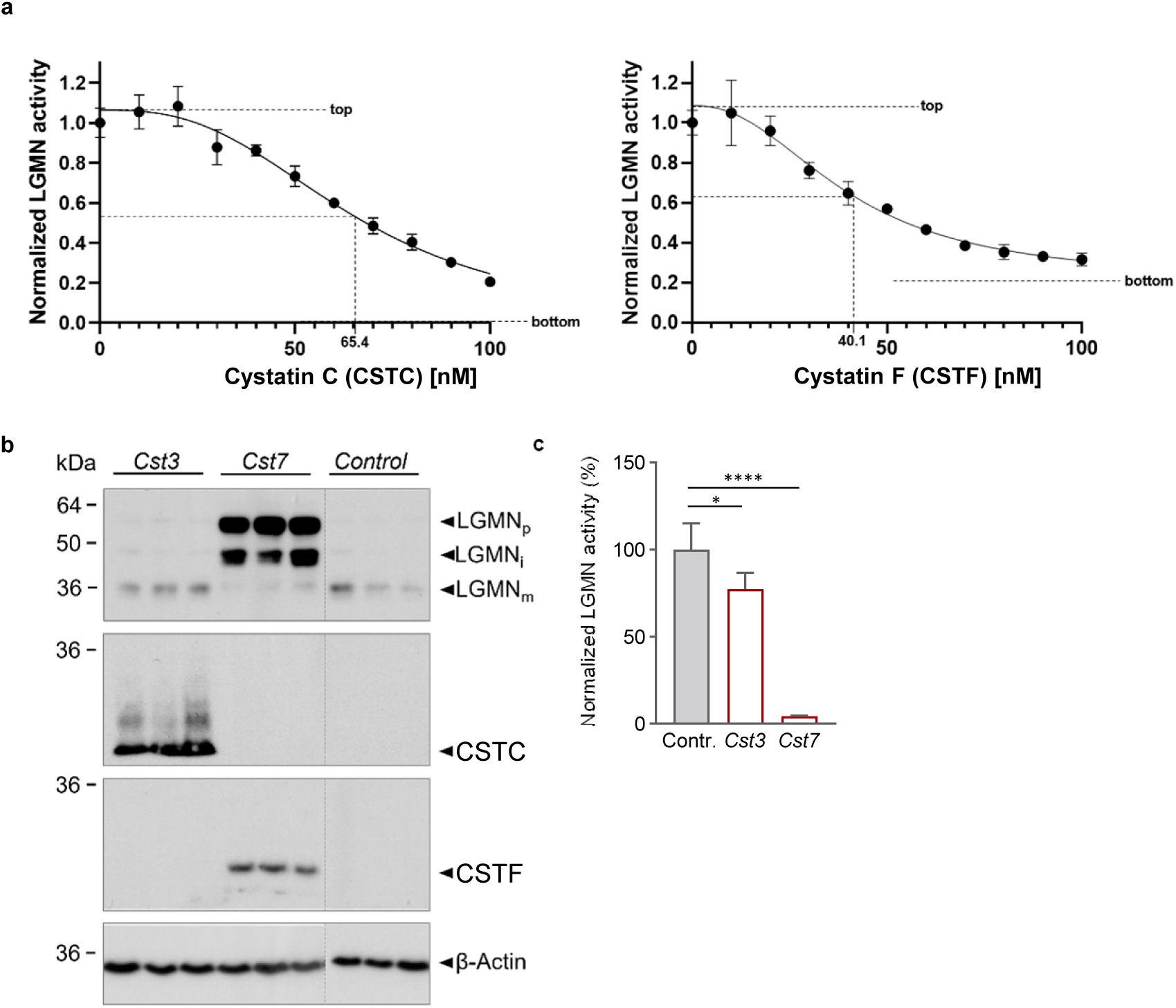
LGMN inhibition by cystatin C and cystatin F. **a,** Dose-dependent inhibition of recombinant LGMN via recombinant cystatin C (CSTC) (left) and cystatin F (CSTF) (right). Nonlinear regression (curve fit; variable slope (4 parameters)), top, bottom and IC50 are indicated. Data points show the mean ± s.d. normalized to LGMN activity at 0 nM inhibitor (N=3 technical replicates). **b**-**c**, HEK 293T cells were transfected with *Cst3*, *Cst7* or pcDNA3 (Control). **b,** Immunoblotting indicates proform (LGMNp), intermediate (LGMNi) and mature form (LGMNm) of LGMN, and CSTC and CSTF expression. β-Actin was probed to verify equal loading. **c,** In vitro LGMN activity assay of cell lysates. Data are mean ± s.d. of biologically independent experiments normalized to control (n=3). Statistical significance was determined by one-way ANOVA and Dunnett’s multiple comparisons post-hoc test: *p<0.05, ****p<0.0001.

**Extended Data Fig. 6:**
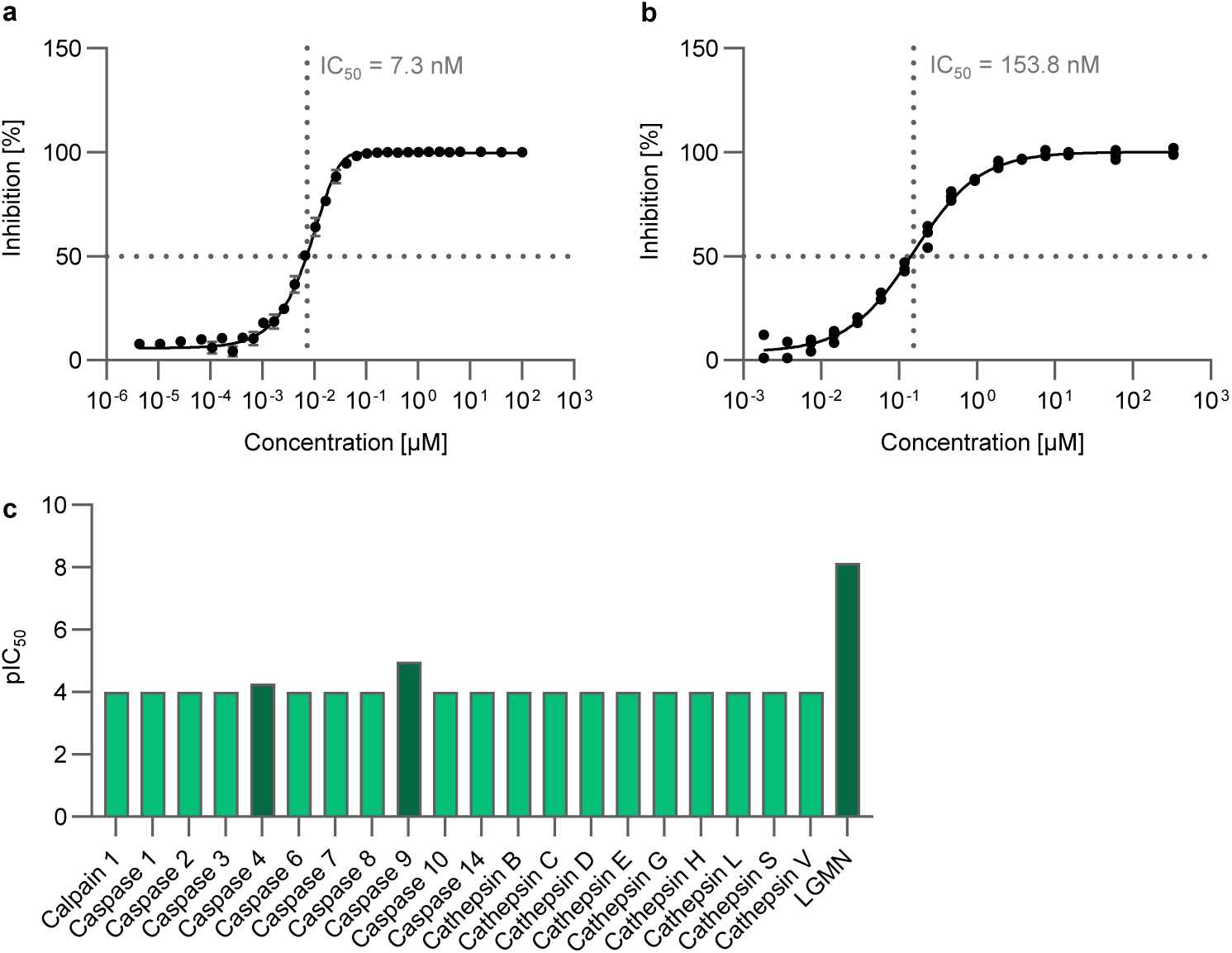
Potency and selectivity of LGMN inhibitor. **a**, Dose-dependent inhibition of pre-activated recombinant LGMN by the LGMN inhibitor. An IC50 of 7.3 nM with a 95% confidence interval from 6.9-7.7 nM was determined (Nonlinear regression curve fit; variable slope (4 parameters)). Data points show the mean of six technical replicates ± s.d. **b**, Dose-dependent inhibition of LGMN in HEK293A cells by the LGMN inhibitor was determined using a cell permeable fluorogenic LGMN substrate. Effects of test compounds were normalized to effect of reference compound and DMSO control. An IC50 of 153.8 nM with a 95% confidence interval from 138.8-170.1 nM was determined (Nonlinear regression curve fit; variable slope (4 parameters)). Data points show the mean of four biological replicates ± s.d**. c**, *In vitro s*electivity screen for the LGMN inhibitor. Proteases with a pIC50 below the limit of detection of 4 are depicted in light green, proteases having a pIC50 >4 in dark green.

**Extended Data Fig. 7:**
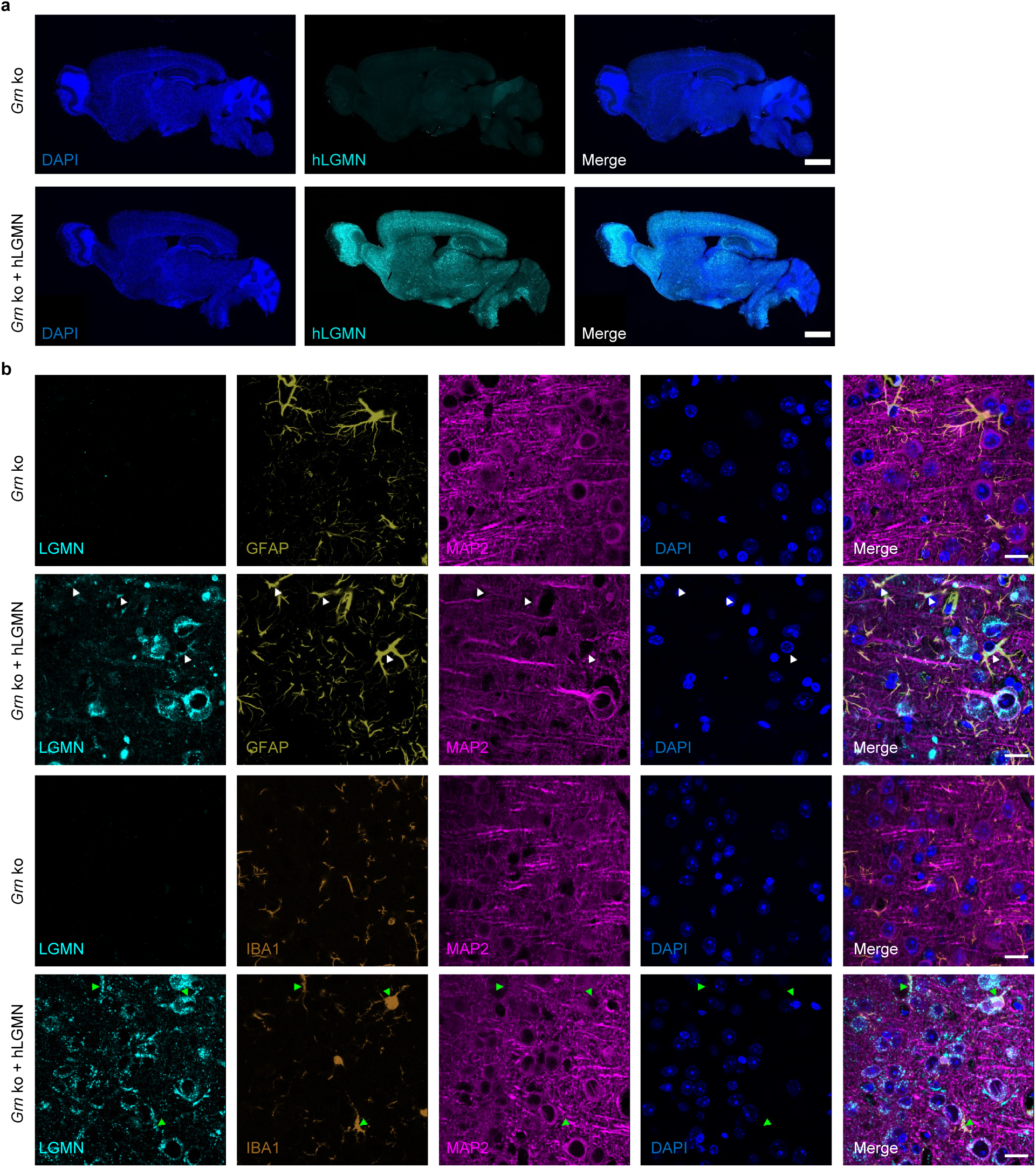
Overexpressed hLGMN localizes predominantly to neurons, astrocytes and microglia. **a**, Immunofluorescence staining of sagittal brain sections 3 months post tail vein injection of AAV-PHP.eB.CAG-hLGMN (DAPI in blue, hLGMN in cyan, n=3 mice, scale bar = 1500 μm). **b**, Immunofluorescence staining for neuronal marker MAP2 (magenta), microglial marker IBA1 (orange) or astrocytic marker GFAP (yellow) and DAPI (blue). White arrowheads show co-localization of hLGMN with astrocytes, green arrowheads co-localization of hLGMN with microglia. (n=3 mice, scale bar = 20 μm).

**Extended Data Table 1:**
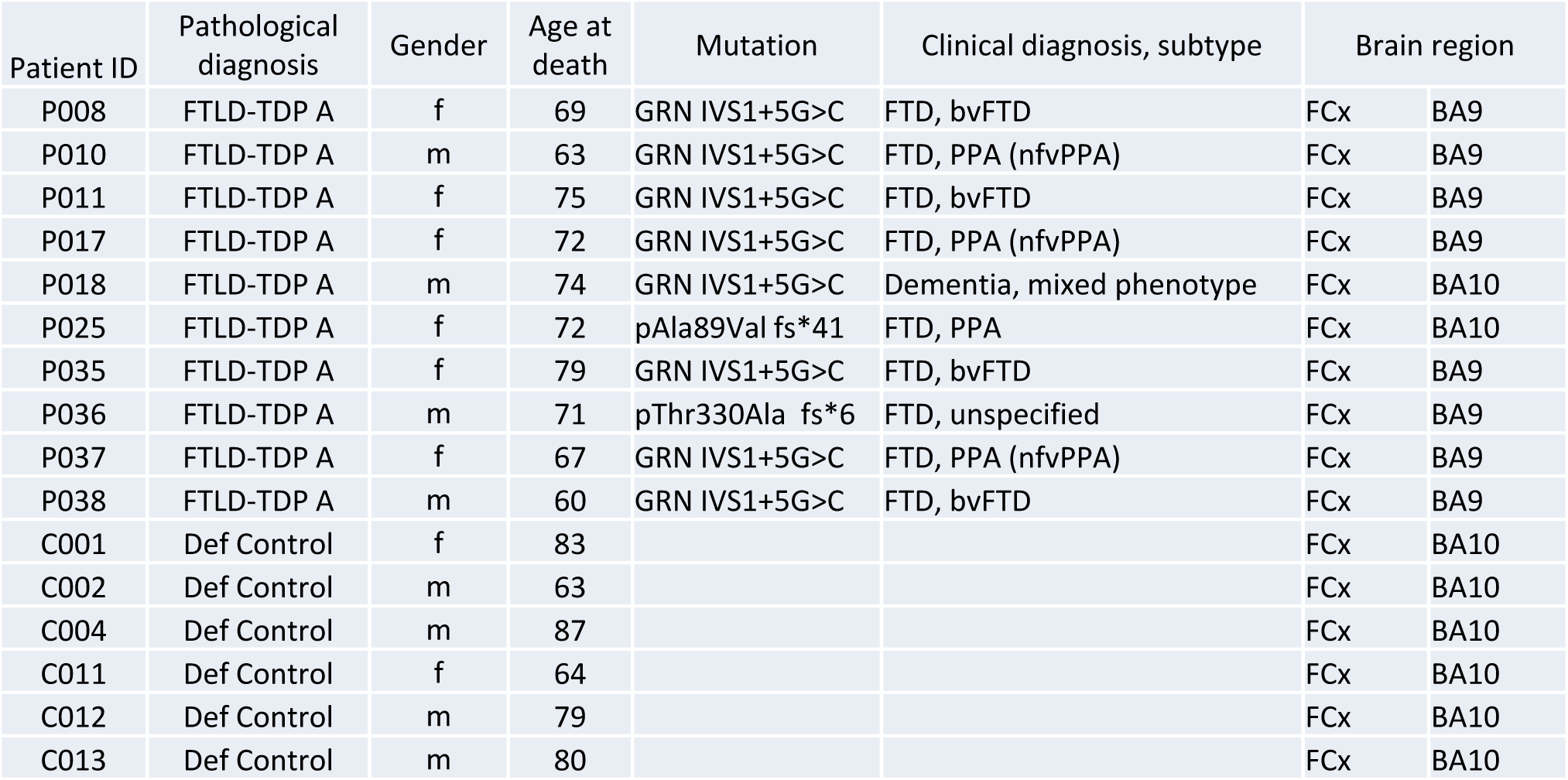
Information on human brain tissue. Summary of clinical, pathological and genetic information of human brain tissue of FTLD-*GRN* and control cases. VIB Department of Molecular Genetics and Antwerp Brain Bank, Institute Born-Bunge: Antwerp, Belgium; m, male; f, female; FCx, frontal cortex.

| Patient ID | Pathological diagnosis | Gender | Age at death | Mutation | Clinical diagnosis, subtype | Brain region |  |
| --- | --- | --- | --- | --- | --- | --- | --- |
| P008 | FTLD-TDP A | f | 69 | GRN IVS1+5G>C | FTD, bvFTD | FCx | BA9 |
| P010 | FTLD-TDP A | m | 63 | GRN IVS1+5G>C | FTD, PPA (nfvPPA) | FCx | BA9 |
| P011 | FTLD-TDP A | f | 75 | GRN IVS1+5G>C | FTD, bvFTD | FCx | BA9 |
| P017 | FTLD-TDP A | f | 72 | GRN IVS1+5G>C | FTD, PPA (nfvPPA) | FCx | BA9 |
| P018 | FTLD-TDP A | m | 74 | GRN IVS1+5G>C | Dementia, mixed phenotype | FCx | BA10 |
| P025 | FTLD-TDP A | f | 72 | pAla89Val fs*41 | FTD, PPA | FCx | BA10 |
| P035 | FTLD-TDP A | f | 79 | GRN IVS1+5G>C | FTD, bvFTD | FCx | BA9 |
| P036 | FTLD-TDP A | m | 71 | pThr330Ala fs*6 | FTD, unspecified | FCx | BA9 |
| P037 | FTLD-TDP A | f | 67 | GRN IVS1+5G>C | FTD, PPA (nfvPPA) | FCx | BA9 |
| P038 | FTLD-TDP A | m | 60 | GRN IVS1+5G>C | FTD, bvFTD | FCx | BA9 |
| C001 | Def Control | f | 83 |  |  | FCx | BA10 |
| C002 | Def Control | m | 63 |  |  | FCx | BA10 |
| C004 | Def Control | m | 87 |  |  | FCx | BA10 |
| C011 | Def Control | f | 64 |  |  | FCx | BA10 |
| C012 | Def Control | m | 79 |  |  | FCx | BA10 |
| C013 | Def Control | m | 80 |  |  | FCx | BA10 |

## Extended data methods

### Experimental mouse models

The second *Grn* ko mouse model and the *Grn* ko/h*TfR*mu/hu ki were previously generated and characterized. *Grn* ko on a C57Bl6/J background were obtained from Jackson Laboratories (JAX strain 013175)^1^. *TfR*mu/hu ki mice (also on a C57Bl6/J background) expressing a chimeric TfR receptor (human TfR apical domain knocked into the mouse receptor) were developed by generating a knock-in of the human apical TfR mouse line using CRISPR, as described previously^2^. Homozygous *TfR*mu/hu male mice were bred to female *Grn*^+/–^ mice to generate *Grn*^−/–^ x *TfR*mu/hu mice. Constitutive *Lgmn* ko mice was generated on a C57Bl6/J background by CRISPR/Cas9 induced deletion of exon 3 of the *Lgmn* gene (842bp) leading to a frameshift knockout by an early stop codon.

### Mouse handling and tissue collection at Denali Therapeutics

Animal weights were collected at the start and end of the study. Mice received therapeutic treatment *via* intravenous (IV) tail vein injection (∼200 µl total injection volume). At indicated endpoints animals were anesthetized with tribromoethanol. Following plasma collection, mice were transcardially perfused with ice-cold PBS. Tissues were then collected, weighed, frozen on dry ice and stored at ™80°C for subsequent analysis.

### Isolation of liver lysosome

As described before^3^ mice were injected with 4 µl/g bodyweight with 17% (w/v) tyloxapol in 0.9% NaCl four days prior to killing and organ removal. After the removal of the liver, the liver was homogenized in 5 ml ice-cold 0.25 M sucrose with four strokes at 1,000 rpm in a Potter-Elvehjem homogenizer. The homogenate was subsequently centrifuged for 10 min at 1,000 x g in a tabletop centrifuge, the supernatant was removed and the pellet re-extracted with another 2 ml of 0.25 M sucrose and centrifuged again. The pooled supernatants (postnuclear supernatant; PNS) were used for differential ultracentrifugation: 9 ml of the PNS were centrifuged at 56,000 x g for 7 min (70.1 Ti rotor, Beckmann, Coulter). The supernatant was removed, the pellet (mitochondria / lysosome fraction; ML) homogenized in 8 ml 0.25 M sucrose and centrifuged again at 56,000 x g for 7 min. The final pellet was resuspended in 3 ml of sucrose solution with a density of 1.21 g/ml. This fraction was overlaid with sucrose solutions with a density of 1.15 g/ml (3 ml), 1.14 g/ml (3 ml) and finally 1.06 g/ml to obtain a discontinuous sucrose gradient. The sucrose gradient was centrifuged at 110,000 x g for 150 min in a swinging bucket rotor (SW41 Ti rotor, Beckmann Coulter). Lysosomes were collected from the interphase between the sucrose solution with a density of 1.14 g/ml and 1.06 g/ml.

### Treatment of mice with PTV:PGRN

PTV:PGRN was generated at DENALI Therapeuticas as previously described (Logan et al.). Mice received 10 mg/kg PTV:PGRN *via* intravenous (IV) tail vein injection (∼200 µl total injection volume) and tissue collected either 7 or 14 days after treatment.

### Transfection of MEF with *Grn* cDNA

MEF *Grn* ko were stably transfected with mouse *Grn* cDNA as described before^4^.

### Cloning of *Cst3* and *Cst7* constructs

Mouse cystation-3 (# MR225378, OriGene) and cystatin-7 (# MR222507, OriGene) myc-tagged cDNA clones were subcloned into the BamHI/EcoRI sites of pcDNA3 using the following primers:

*Cst3* (BamHI) forward 5’-CCCGGATCCCCAACCATGGCCAGCCCGCTGC-3’ *Cst7* (BamHI) forward 5’- CCCGGATCCCCAGCCATGCCCTGGTCCTGG ™3’ *Cst3/7* (EcoRI) reverse 5’- CGGAATTCTTAAACCTTATCGTCGTCATCC ™3’ Six hundred three base pairs (603 bp) encoding for cystatin-7 were cloned into the BamHI/EcoRI sites of pAAV-hSyn-EGFP (# 50465, Addgene). Serotype AAV9 (pAAV-GOI) virus particles were produced in HEK 293T cells by co-transfection with pAAV-RC, pHelper and pAAV-GOI using Lipofectamine 2000 (ThermoFisher Scientific). After 72 h cells were detached with 0.5 M EDTA and centrifuged for 5 min at 3000 rpm. The pellet was resuspended in 250 µl Neurobasal Medium (ThermoFisher Scientific) followed by 4 freeze-thaw cycles in a dry ice ethanol bath and a 37°C water bath for 2 min per bath and cycle. Centrifugation for 10 min at 10,000 x g generated the viral particles containing supernatant. AAV was stored at ™80°C until further use.

### *In vitro* determination of pIC50 for the LGMN inhibitor

A modified fluorescence-based activity assay was used to assess the potency of the LGMN inhibitor to inhibit LGMN activity *in vitro*. Recombinant LGMN stored in 25 mM HEPES/NaOH pH 7, 300 mM NaCl, 200 mM trehalose was activated at a concentration of 0.1 mg/mL for 2 h at room temperature in 50 mM sodium actetate pH 4.0. Assay buffer was 20 mM Na-phosphate/citrate pH 5.8, 1 mM EDTA, 0.1% CHAPS. The pre-activated LGMN (c=0,1 nM) was incubated with10 µM of peptide Z-AAN-Rh110 and different doses of LGMN inhibitor. Excitation and emission wavelengths were 485 nm and 520 nm, respectively. Fluorescence was measured after 15 min and 30 min incubation.

### Determination of pIC50 for the LGMN inhibitor in HEK293A cells

HEK293A overexpressing LGMN were cultured using DMEM (Gibco 31966) supplemented with 10% FCS, 1% Penicillin/Streptomycin and 0.8 mg/ml Geneticin. On day before the assay 16.000 cells/well were plated in 384 well plates. At the assay day cells were treated with dose responses of LGMN inhibitor or DMSO control. Final DMSO concentration for all conditions was 0.6%, and compounds were tested in a 16- point-dose response starting at 60 µM. Cells were incubated with LGMN inhibitor for 2 h, prior to addition of fluorogenic substrate **S** (Roche, final concentration 60 µM)^5^. After addition of substrate cells were incubated for 5 h and the effect was detected by measuring fluorescence intensity at 460 nm after excitation at 380 nm in the living cells.

### Protease selectivity screen

To test the selectivity of the LGMN inhibitor, pIC50 values for different proteases were determined using the Protease Assay Service from REACTION BIOLOGY (https://www.reactionbiology.com/services/target-specific-assays/protease-assays). pIC50 for LGMN was determined by Roche, Basel.

### Immunofluorescence of mouse brain tissue

Freshly isolated brain hemispheres were post-fixed in 4% paraformaldehyde in PBS (Thermo Scientific, J19943-K2) for 24 h at 4°C. Brain hemispheres were transferred to PBS with 0.02% NaN3 and stored at 4°C until further use. Brain hemispheres were cut in 40 µm sections using a vibratome (Leica VT1200S). Immunofluorescence was performed on free floating sections. Sections were blocked with blocking solution (4% bovine serum albumin (BSA, Sigma A8022-100G) and 0.1% fish skin gelatin (Sigma G7041) in PBS with 0.4% Triton-X) for 1 h at RT. Primary antibodies, hLGMN (R&D, AF 2199, c= 0.4 µg/ml), MAP2 (Novus Biologicals, 1:1000, NB300-213), IBA1 (GeneTex, 10042, c=0.06 ng/µl), GFAP (DAKO, Z0334, c=2.9 µg/ml) were diluted at the indicated concentration in blocking buffer and brain sections were incubated over night at 4°C in the primary antibody solutions. The next day the sections were washed three times with PBS with 0.1% Tween® 20 and subsequently incubated with fluorophore-coupled secondary antibodies (Thermo Fisher Scientific, donkey, c=4 µg/ml) diluted in secondary antibody buffer (4% bovine serum albumin (BSA, Sigma A8022-100G) and 0,1% fish skin gelatin (Sigma G7041) in PBS) with DAPI (c=1 µg/ml) for 2 h at RT. After three washing steps in PBS with 0.1% Tween 20 sections were mounted with Fluoromount-GTM (Invitrogen, 00-498-02).

### Image acquisition

Full brain hemispheres were imaged at 4x with a Leica DMi-8 epifluorescence microscope with a DFC9000GT camera. Image mosaics were merged using the Leica LAS X software suite. Representative images were acquired on a Zeiss LSM800 Axio Observer 7 confocal microscope with a 40x oil immersion objective (NA 1.3). A z-stack was acquired and confocal images show an orthogonal projection of six images with a z-stack size of 0.4 µm. Post-processing (adjustment of brightness) of full brain scans was performed in ImageJ and brightness of confocal images was adjusted with the Zen 3.2 (blue edition) software in the same way for all conditions.

### Additional antibodies used in figures

| Antibodies | Dilution |  | Source |
| --- | --- | --- | --- |
|  | IB | IF |  |
| Goat anti-mouse CatD | 1:1,000 | -- | # sc-6486, Santa Cruz |
| Goat anti-mouse CatL | 1:1,000 | -- | # AF965, R&D Systems |
| Goat anti-cystatin-3 | 1:1,000 | -- | # AF1238, R&D Systems |
| Rabbit anti-GFAP | -- | 1:1,000 | # Z0334, DAKO |
| Rabbit anti-mouse GFAP | 1:5,000 | -- | # GA52461-2, DAKO |
| Rabbit anti-mouse IBA1 | 1:1,000 | -- | # GTX100042, GeneTex |
| Rat anti-mouse LAMP1 | 1:1,000 | -- | # sc-19992, Santa Cruz |
| Rabbit anti-mouse tubulin β3 (TUJ) | 1:1,000 | -- | # 5568S, Cell Signaling |

